# Integrative spatial and multi-omic profiling in bladder cancer links L1 retrotransposition to extrachromosomal DNA, genomic instability, and viral mimicry response

**DOI:** 10.1101/2025.07.30.667694

**Authors:** Sophia J. Pribus, Ivana Osredek, Jan Otonicar, Milena Simovic-Lorenz, Michael Scherer, Sergio Manzano-Sanchez, Andreas Kienzle, Urja Parekh, Vladimir Benes, Pooja Sant, Philipp Mallm, Karsten Brand, Angelika B. Riemer, Holger Sültmann, Christoph Plass, Mladen Stankovic, Jan O. Korbel, Tobias Rausch, Aurélie Ernst

## Abstract

Bladder cancer is one of the most frequent cancers and shows high recurrence rates. Despite recent advances, key knowledge gaps remain in understanding the molecular mechanisms of disease progression, which would support the development of early detection methods and effective personalized treatments. We apply integrated multi-omics and spatial analyses in a cohort of 49 bladder cancer patients to comprehensively profile genetic, epigenetic, transcriptomic, and spatial features of bladder cancer, alongside cell-free DNA blood analysis. Combining low-pass whole-genome cell-free DNA sequencing, Oxford Nanopore long-read tumor DNA sequencing, RNA-sequencing, and spatial transcriptomics, we provide insights into molecular alterations driving bladder cancer. We show frequent somatic LINE-1 (L1) insertions, with up to more than 500 insertions per tumor. We find that L1 insertions are active and occur early in bladder cancer development. We link aberrant somatic L1 insertion in bladder cancer with downstream genomic rearrangements and chromosomal instability, with an excess of structural variants and extrachromosomal DNA (ecDNA) in patients with particularly high L1 counts. By detecting ecDNA within tissue architecture using spatial transcriptomics, we identify the localization of ecDNA to distinct spatial clusters with differential expression of *APOBEC3B* and immune response pathways. These results, combined with replication timing analysis and gene set enrichment analysis (GSEA), offer evidence for the previously hypothesized viral mimicry response to L1 retrotransposition, mediated via APOBEC3B-editing, the cGAS-STING pathway, and RIG-I and MDA5 responses.

## INTRODUCTION

Bladder cancer is one of the most frequent malignancies worldwide, with an estimated 573,000 new cases and 213,000 deaths occurring globally each year^1^. It arises from the urothelial lining of the bladder and is classified into two major types: non-muscle invasive bladder cancer (NMIBC) and muscle invasive bladder cancer (MIBC)^2^. NMIBC accounts for approximately 70-75% of cases at diagnosis and when diagnosed early shows high rates of survival^3,4^; MIBC is more aggressive and has a worse prognosis^2,4^. Despite improvements in detection and treatment, bladder cancer remains challenging to treat due to frequent transitions from NMIBC to MIBC^5^, propensity for recurrence and metastasis^2^, and resistance to current therapies^2,6–10^. These challenges are especially pronounced in the surveillance of recurrent disease^2^, where clinical ambiguity can lead to overtreatment or delayed intervention. Overall, there is a lack of understanding of the clinical and genomic features that influence disease aggressiveness; addressing this gap would support the development of improved diagnostic and prognostic tools, and facilitate the development of more effective treatment strategies. Thus, a more comprehensive molecular characterization of bladder cancer development may pave the way to significantly improve patient outcomes.

Earlier detection and improved risk stratification have the potential to significantly decrease bladder cancer-related death^11–13^. While cystoscopy and urine cytology are widely used in clinical practice, these methods are invasive, costly, and often fail to detect tumors at early stages^14^. Moreover, they lack specificity and the sensitivity to distinguish between benign and malignant lesions, or to predict which patients are at risk of disease progression^2^. A deeper understanding of the drivers of disease initiation and development through integrated clinical and genomic profiling has the potential to reduce invasiveness and improve accuracy of early detection and risk stratification for patients. In addition, the lack of comprehensive understanding of the disease hinders the development of targeted treatment approaches. While a number of molecular studies^15–19^ of bladder cancer have identified key driver genetic alterations and pathways, none to our knowledge have combined comprehensive genomic, epigenomic, transcriptomic, and spatial data to reveal novel, bladder cancer-specific disease mechanisms. Such a study could lay the basis to understand recurrence mechanisms, identify innovative therapeutic targets, and explore candidate biomarkers for personalized interventions.

The potential of advanced genomic technologies, including long-read sequencing and spatial transcriptomics, to improve our mechanistic understanding of bladder cancer progression has, to our knowledge, not been fully exploited. One such mechanism that can be explored through the use of such next-generation sequencing technologies is the somatic retrotransposition of LINE-1 (L1). Current estimates suggest that L1 sequences make up roughly one-fifth of the human genome, and while the majority of these are inactivated via epigenetic silencing, the small fraction of active L1s are hypothesized to have an impact on cancer progression^20^. This has been suggested in a bladder cancer context, where L1 elements become active via hypomethylation and contribute to genomic instability^21^. Increasing evidence indicates that the DNA damage associated with L1 insertion may influence downstream mechanisms of genomic alterations and immune response via viral mimicry^20^. However, comprehensive studies of these mechanisms are lacking in clinical samples, including in bladder cancer. Linking L1 insertions with aggressive disease characteristics such as genomic rearrangements, amplifications, ecDNA and upregulated immune response would solidify the potential of L1 as a novel bladder cancer biomarker.

In cell lines, L1 insertions have been shown to upregulate DNA damage mechanisms by causing double-stranded DNA breaks and have been hypothesized to contribute to chromosomal instability in cancer^20,22,23^. Importantly, pan-cancer analyses have shown that somatic L1 retrotransposition induces breakage-fusion-bridge cycles and chromothripsis^20^, known precursors to extrachromosomal DNA (ecDNA)^24^; however, a link between L1 insertions and ecDNA presence has not been established in a cancer context. This is especially interesting, given that L1 insertions have been shown to trigger APOBEC3B-mediated mutagenesis^20^, and APOBEC3B activity has been shown to facilitate ecDNA evolution^25^. Demonstrating a link between L1 insertion count and ecDNA presence could suggest a potential “double-hit” mechanism via which L1 insertions concurrently trigger APOBEC3B and chromosomal instability leading to ecDNA formation, and maintained APOBEC3B activity further mutates ecDNA, facilitating downstream tumor aggressiveness.

In cell lines and in tumors outside the bladder cancer context, L1 expression has been linked to immune response via “viral mimicry”, a process by which endogenous virus response mechanisms are activated by non-viral genetic material in the cytoplasm. It has been proposed that L1 insertions are a source for double-stranded DNA (dsDNA), double-stranded RNA (dsRNA), and DNA:RNA hybrids, all of which, when detected in the cytoplasm, may elicit the innate type I interferon (IFN-I) response^20,26,27^. To substantiate these claims, it is important to first establish the upregulation of such response pathways, including the cGAS-STING pathway (which responds to dsDNA and DNA:RNA hybrids) and the RIG-I and MDA5 sensors (which respond to dsRNA). This can be addressed through both RNA expression analysis and spatial transcriptomics for additional spatial resolution of response pathways.

By integrating multi-omics and spatial approaches, we aimed to gain a comprehensive view of the genetic, epigenetic, transcriptomic, and spatial characteristics of bladder tumors, alongside blood-based analysis of cfDNA. We included both NMIBC and MIBC patients across tumor grades and stages to capture the entire spectrum of bladder cancer. The combination of low-pass whole-genome sequencing of cfDNA, Oxford Nanopore long-read sequencing for tumor genomes, matched germline DNA sequencing, RNA-sequencing, and spatial transcriptome analysis provides a holistic view of the (epi)genetic alterations and gene expression changes driving bladder cancer. In particular, we identify for the first time in a clinical bladder cancer cohort a comprehensive connection between L1 insertions, chromosomal rearrangements including ecDNA, and potential downstream viral mimicry effects. By generating comprehensive molecular insights into bladder cancer biology, this work suggests the potential of L1 elements as a novel biomarker in bladder cancer and elucidates possible mechanisms of disease progression via L1 insertion-mediated genomic instability.

## RESULTS

### Patient cohort

Understanding the molecular landscape of MIBC and NMIBC is essential to improve risk stratification and to develop early detection approaches as well as targeted treatment strategies. We present a comprehensive multi-omic characterization of tumors and blood from 49 patients with MIBC, NMIBC or benign bladder lesions (**Figure 1a**). The tumors represent the whole disease spectrum of bladder cancer, including low-grade as well as high-grade tumors of all stages. A full description of the cohort is displayed in **Supplementary Table A**. We analyzed blood cell-free DNA (cfDNA) using low-pass whole-genome sequencing (n=49, mean coverage: 4.5x). We performed long-read whole-genome sequencing using Oxford Nanopore technology (ONT) (n=23, mean coverage: 32x, N50 read length: 15.7Kbp) and short-read sequencing of matched germline DNA (n=23, mean coverage: 17x). Furthermore, we conducted spatial transcriptomics analyses using the Visium platform for 10 tumors representative from the different stages and grades, and total RNA-sequencing (RNA-seq) for 17 tumors (see **Supplementary Table A** for details).

**Figure 1.**
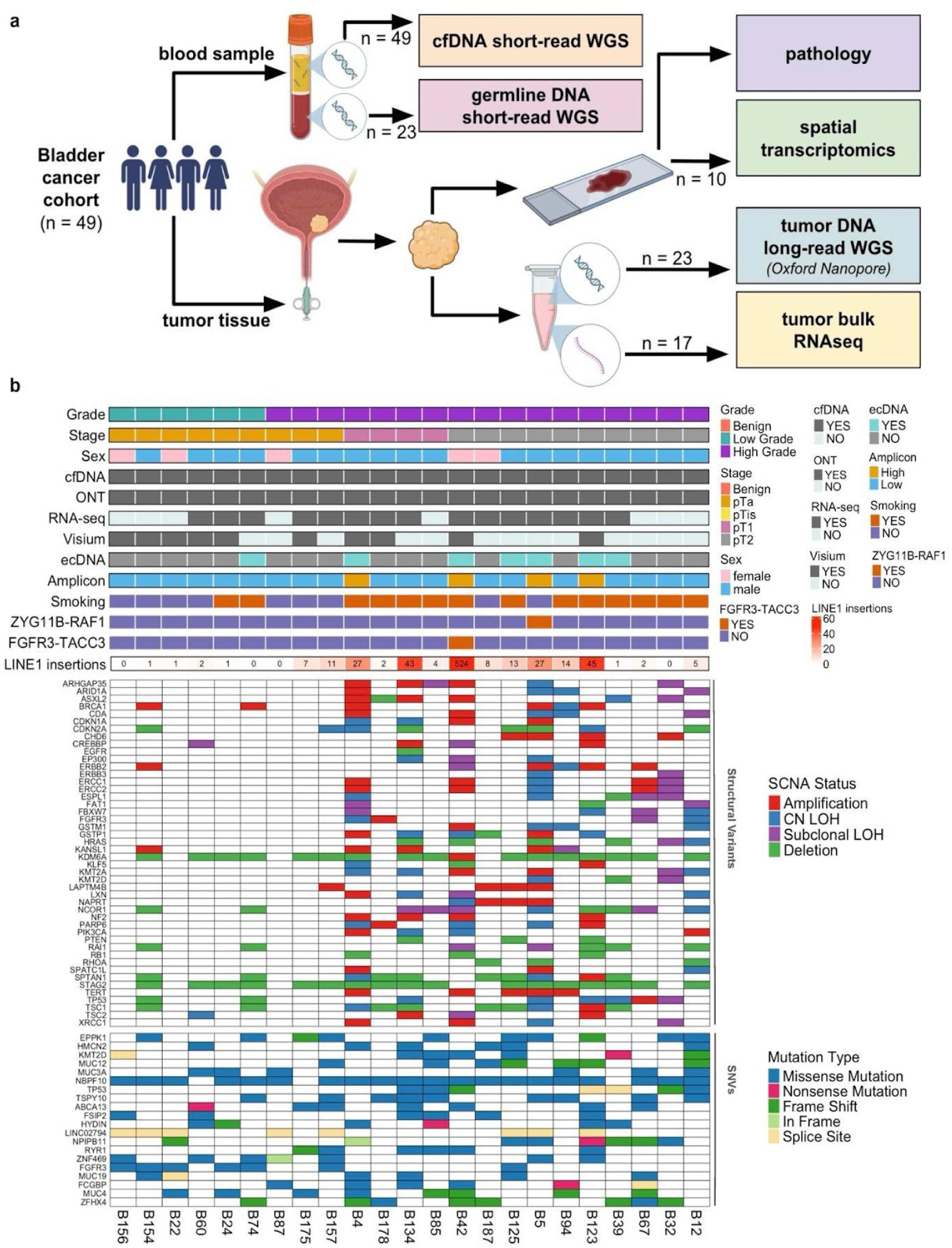
Multi-omic analysis reveals multiple forms of genome instability in bladder cancer. **a)** Schematic of patient cohort and data types (created with BioRender). 49 bladder cancer patients underwent blood draws and transurethral resection of bladder tumor. Blood samples were used for cfDNA analysis with short-read WGS. Tumor samples were used for spatial transcriptomics, genomic (long-read WGS) and RNA-Seq analysis. **b)** Oncoplot summarizing patient-specific tumor characteristics (stage and grade, ecDNA status, somatic L1 insertions), sex, available data types, driver gene-fusions and other alterations in key bladder cancer driver genes detected by long-read sequencing. CN = copy number, CN LOH = copy-neutral loss of heterozygosity.

We first explored gene-level alterations that could contribute to cancer development in our cohort. Analysis of somatic copy-number alterations (SCNAs) revealed recurrent somatic alterations affecting established bladder cancer driver genes (**Figure 1b**). The most frequently altered genes included *KDM6A* (altered in 73.9% of the tumors, with 94.1% showing deletions), *STAG2* (73.9%, 100% deletions), *NCOR1* (43.5% with deletions and subclonal loss of heterozygosity (LOH), each contributing 40%), *SPTAN1* (43.5%, 70% of deletions), *TP53* (43.5%, 40% LOH), *TSC1* (43.5%, 70% of deletions) and *RAI1* (39.1%, 55.6% of deletions). We also analysed single nucleotide variants (SNVs) (**Figure 1b)**. The most frequently altered gene was *NBPF10*, mutated in 87% of the tumors through missense mutations (100%). Other commonly affected genes included *EPPK1* (43.5%, 80% missense), *TSPY10* (39.1%, 100% missense), and *LINC02794* (34.8%, 100% splice site mutations). A full list of affected genes and the respective alterations are available in **Supplementary Tables B-C.**

We further identified 65 high-confidence gene fusions, supported by ONT reads and RNA-seq data (**Supplementary Table D**). The *FGFR3-TACC3* fusion (**Supplementary Figure 1a**), previously reported in bladder cancer^28,29^, leads to constitutive activation of FGFR3 signaling, promoting cell proliferation and survival. The *ZYG11B-RAF1* fusion (**Supplementary Figure 1b**), not previously identified in bladder cancer, potentially drives cancer development, as suggested by gene fusions involving *RAF1* identified in other cancers and by activations of *RAF1* through fusion-independent mechanisms such as amplification in approximately 12–20% of MIBC. In the germline, we also identified 12 high-impact pathogenic variants using Variant Effect Predictor (VEP)^30^, 33 variants of moderate impact and two modifier variants, respectively (see **Supplementary text** for details).

### Long-read, spatial, and cell-free DNA analysis enable detailed dissection of genomic instability and retrotransposition events within the tissue context

We next extended our analysis beyond the gene level by applying an integrative multi-omic profiling approach to uncover mechanisms driving cancer evolution and crosstalk with the microenvironment. For all samples in the cohort, we dissected the interplay between genome and epigenome from ONT sequencing data as well as cfDNA analysis. We further explored these relationships spatially, mapping the genetic subclone composition and spatial patterns within the tissue architecture. To demonstrate the value of this multi-omic approach, we highlight our comprehensive analysis of tumor, cfDNA and germline DNA from patient B123 in **Figure 2**.

**Figure 2.**
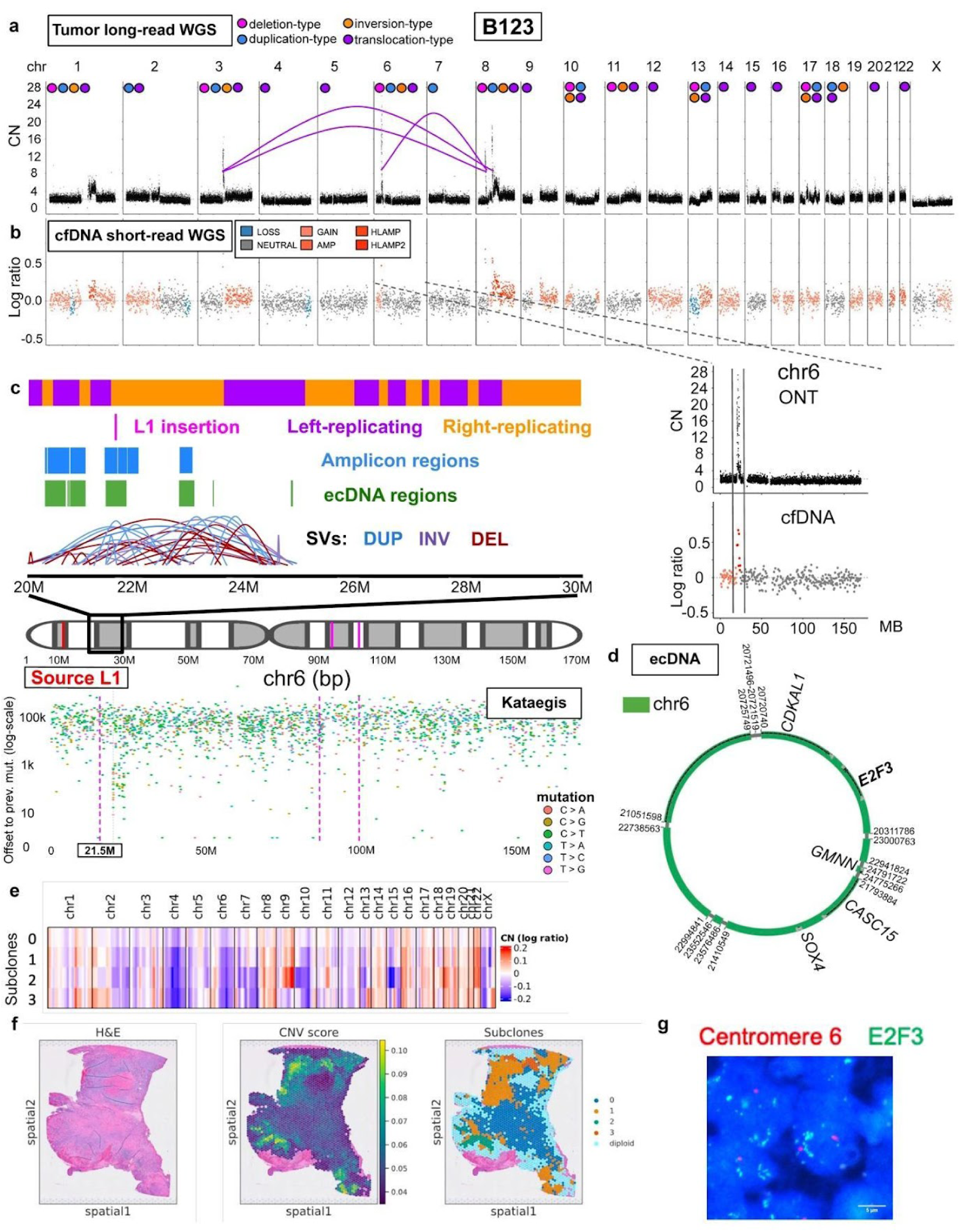
Long-read, spatial, and cell-free DNA analysis allows dissecting the landscape of tumor cell clones, extrachromosomal circular DNA and retrotransposition within the tissue architecture. **a)** Long-read WGS of B123 tumor tissue reveals broad copy-number variation and sporadic high copy-number amplifications. Chromosomes are labeled with a subset of the detected structural variation events; a subset of interchromosomal translocation events where both ends border amplicon events (within 100kbp) are displayed as purple arcs. **b)** cfDNA short-read WGS shows matched copy-number variations between B123 blood and tumor samples and in particular, specific amplification events on chromosomes 3, 6, and 8. LOSS: CN < 2, NEUTRAL: CN = 2, GAIN: CN = 3, AMP: CN = 4, HLAMP: CN = 5, HLAMP2: CN >= 6. **c)** Schematic of focal genomic events on 6p21 in sample B123, including L1 insertions, amplicon regions, ecDNA regions, structural variations, and kataegis events. **d**) Schematic of ecDNA detected in B123, including amplifications of *E2F3*, *CDKAL1*, *GMNN*, *CASC15*, and *SOX4* (chromosome schematic from BioRender). **e**) Copy-number inference from Visium identifies genetic clones with distinct copy-number profiles (sample B123). **f)** Hematoxylin and eosin stain (left), CNV score from Visium (middle) and genetic subclones shown in E (right) (sample B123). **g)** FISH validation of ecDNA carrying the *E2F3* oncogene (B123).

In patient B123, ONT sequencing revealed multiple focal amplifications and 137 structural variants (SVs), including 47 inter-chromosomal events (**Figure 2a**). Across the whole cohort, we identified at least one somatic SV in all ONT-sequenced tumors (**Supplementary Figure 2**). For patient B123, copy-number alterations in cfDNA reflected those identified using ONT sequencing in the matched tumor tissue (**Figure 2b**). However, when considering the whole cohort, only a minority of cfDNA samples showed detectable copy-number changes, with evidence for enrichment in high-grade tumors (4/25 high-grade tumors, 0/23 low grade, p=0.06, Fisher’s exact test, one-sided, **Supplementary Figure 3**). There was also evidence for chromothripsis in 12/23 patients (**Supplementary Figure 2**), including patient B123 (**Figure 2a**). We further detected ecDNA in 7/23 patients, harboring oncogenes such as *CCND1*, *MDM2* and *E2F3* (**Supplementary Table A and E**), including B123 (**Figure 2c-d**). Beyond ecDNA, we also detected indications of oncogene activation through enhancer hijacking by computational prediction with *pyjacker*^31^ (**Supplementary Figure 4a-d**), identified as DNA rearrangements juxtaposing an enhancer nearby an oncogene coupled with a strong overexpression. Notably, tumor-promoting genes (e.g., *UCHL1*) putatively activated through enhancer hijacking were largely independent from those activated through ecDNA formation (**Supplementary Figure 4e**). Alignment to the CHM13 (T2T) assembly^32^ and calculation of a chrY expression signature^33^ further revealed loss of chromosome Y (LOY) in 23.5% of male patients (**Supplementary Table A, Supplementary Figure 5a-b**), including patient B123, and additional chromosome Y copy number variations. Importantly, these LOY patients showed significant upregulation of metastasis and tissue invasion pathways (**Supplementary Figure 5c**). Finally, using the matched spatial transcriptome, we performed copy-number (CN) inference, quantified genome-wide CN score and identified up to seven genetic clones per tumor (**Figure 2e-f, Supplementary Figure 6)**.

Interestingly, facilitated by the long ONT reads, we identified a high numbers of somatic Long interspersed element 1 (LINE-1, L1) insertions (19/23 patients with at least one somatic L1 insertion; 750 total L1 insertions across patients). For one sample (B125), short-read and long-read L1 discoveries were directly compared, with two-thirds of the somatic L1 predictions also being confirmed through short-read data (**Methods**). For somatic L1 insertions with confidently identified source elements, we compared their source regions to known L1 source element loci, which identified two novel source L1 elements located on chromosomes 1 and 9, respectively (**Methods**; **Supplementary Table F**). We also found 11 “hot” source L1 regions that either gave rise to multiple somatic L1s within the same patient (B42), or that were active in multiple patients (**Supplementary Table G**). Nine of these “hot” source L1 regions overlapped with previously reported source L1s, whereas two were novel.

In patient B123, we further identified positional proximity (within 10 kbp), but not precise overlap, of an L1 insertion and ecDNA segment breakpoint with a kataegis mutational event (likely *APOBEC3B*-mediated, given the C>T transition), proximal to a replication origin site (**Figure 2c**). We validated somatic L1 insertion activity by immunofluorescence analysis of L1 ORF1p expression and ecDNA structures by FISH and immunofluorescence analysis of oncogene expression on consecutive tissue sections, respectively (**Figure 2g, Supplementary Figure 7-8**). These results led us to further explore the putative role of initiating events that might contribute to L1 mobilisation.

### Microbial infection as a potential driver of L1 mobilization

We next investigated the potential impact of microbial species on bladder cancer initiation and progression. Using the ONT data, we identified *Cutibacterium* as a common microbial species in bladder cancer tissue, as previously described; however, as *Cutibacterium* is frequent on the skin, this may potentially be a contamination (**Supplementary Figure 9a)**. In patient B42, however, we additionally detected reads from *Anaerococcus* (**Supplementary Figure 9b)**, a Gram-positive, anaerobic bacteria that is part of the normal human microbiota, but can become an opportunistic pathogen. *De novo* assembly confirmed the presence of bacterial DNA in the whole-genome sequencing data of this sample, and the matched RNA sample also showed *Anaerococcus* reads. We then analysed 23 additional bladder cancer samples from PCAWG^34^ (BLCA-US) but did not identify any additional tumors with such a high *Anaerococcus* abundance. Even though not a frequent event, *Anaerococcus* infection may contribute to carcinogenesis by inducing reactive oxygen species (ROS) production^35^. These reactive oxygen species (ROS) can cause hypomethylation of L1 elements in bladder cells^36^, potentially triggering the large number of somatic L1 insertions (>500) in B42.

### L1 elements are activated via demethylated promoters and associated with genomic instability, including ecDNAs

We next turned to investigate the regulation of L1 activation and genome integration. We first analysed the ONT-derived DNA methylation data of the patient with the highest number of somatic L1 insertions, namely B42 (**Figure 3a**). One mechanism for L1 activation is via promoter hypomethylation, as previously suggested in the literature^37^. We found that methylation levels were significantly lower at the L1 promoters compared to the bodies of source L1s (p = 4.48e-36, Rank-Biserial correlation (effect size) = 0.207, one-sided Mann–Whitney U test; **Figure 3d**). As a comparison, in patient B60, who showed no L1 activity, the same L1 source loci displayed methylated promoters, supporting the association between promoter hypomethylation and retrotransposition activity (**Supplementary Figure 10a**). Since some L1 insertions can transduce adjacent DNA sequences of the source L1 element to the insertion site, it is sometimes possible to trace the origin of a somatic L1. Using this approach and the long-read data, we performed *de novo* assembly for L1 elements and identified two instances where a somatic L1 served as the source for additional insertions. In one of these cases, the source element in GRCh38 could be clearly traced (**Figure 3c, Supplementary Figure 10e**). These “multi-jumps”, where a somatic L1 insertion induces further L1 retrotransposition, prompted us to investigate the promoters of all full-length somatic L1 insertions, an analysis that revealed low overall methylation levels, supporting the notion that even somatic L1 insertions retain their retrotransposition competence in B42 (**Supplementary Figure 10c, Methods**).

**Figure 3.**
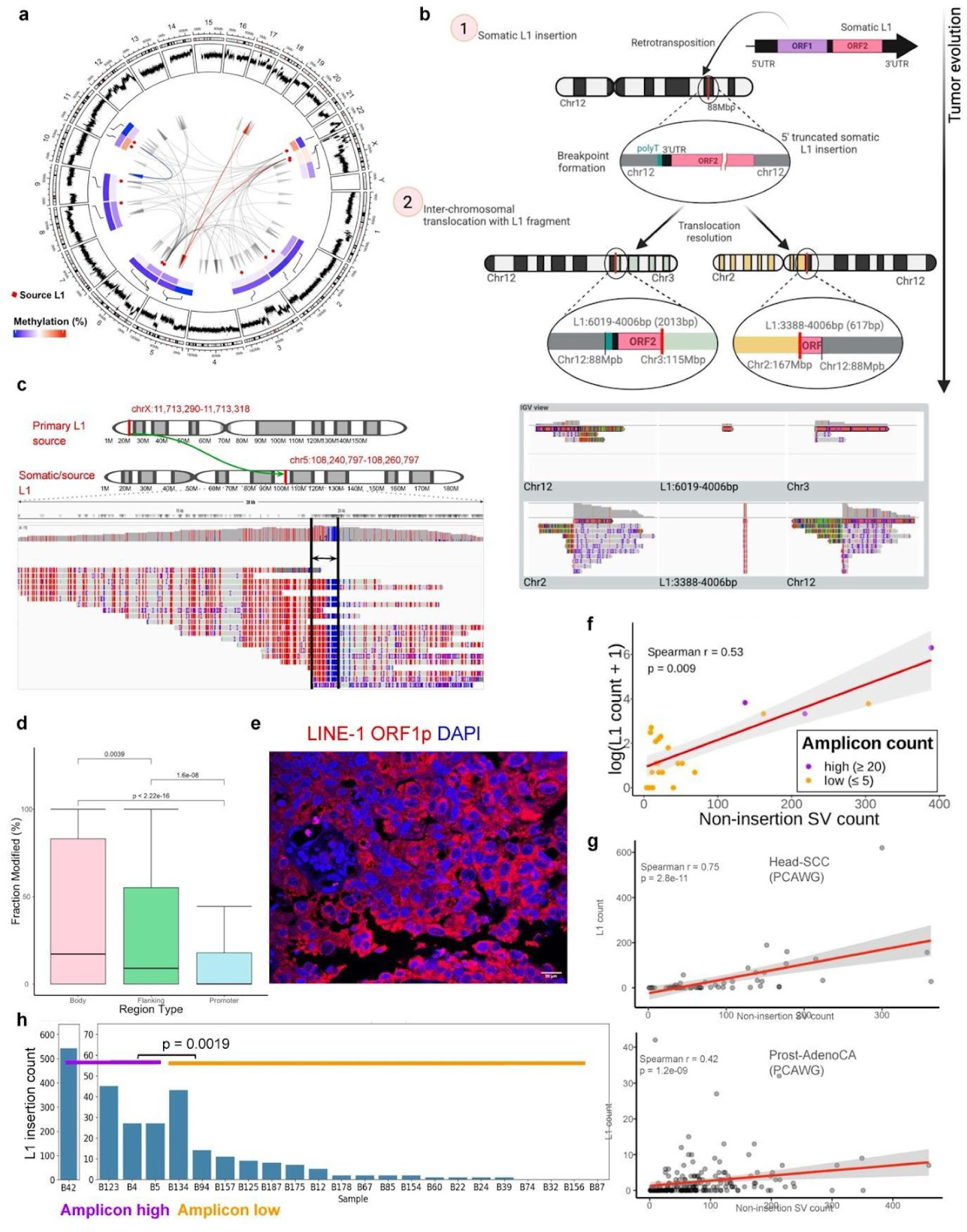
L1 element activation via demethylated promoters and association with genomic instability. **a)** Circos plot of the B42 tumor. The outermost track displays genome-wide copy number alterations. The second track shows methylation levels of source L1 promoter regions and the third track methylation of their corresponding body. The innermost track highlights active L1 elements, with arrows indicating the direction of retrotransposition from the source to the target genomic location. Highlighted arrows in blue and red indicate L1 multi-jumps. **b)** Example of a somatic L1 insertion embedded between somatic structural variant (SV) breakpoints on chromosomes 2, 3 and 12. **c)** Example of a multi-jump L1 event detected in B42, visualized in IGV. The 2-color 5mC IGV mode visualizes unmethylated bases in blue and 5-Methylcytosine (5mC) in red. **d)** Methylation levels of source L1 elements (Mann-Whitney U-test) between promoter (n=1032), body (n=2631), and 500 bp flanking regions (n=442). **e)** Validation of LINE-1 expression by staining for Long interspersed element 1 (LINE-1) open reading frame 1 protein (ORF1p) on matched tissue sections. **f)** Correlation between the number of L1 insertions (log-scaled) and non-insertion SVs across all samples in the bladder cancer cohort (n = 23). Each data point shows one tumor sample. **g)** Correlation between the number of L1 insertions and non-insertion SVs across PCAWG for Head-SCC (n = 57) and Prost-AdenoCA (n = 234). **h)** Counts of L1 elements across tumors. Amplicon-high tumors (n=4) are enriched for L1 insertions versus amplicon-low tumors (n=19) (p = 0.0019, Rank-biserial effect size = 0.947, one-sided Mann-Whitney U-test).

Somatic L1 insertions in B42 appear to be randomly distributed throughout the genome without any apparent clustering (**Supplementary Figure 10d**). However, there was evidence that some somatic L1 elements are directly associated with SV breakpoints, as we identified genomic breakpoints in which L1 fragments are embedded at the junction. These intricate SV events generate complex breakpoint patterns that are not directly mapped by conventional SV and mobile element callers (**Methods**). This was exemplified by an inter-chromosomal translocation of chr2 and chr12 with a 617bp long L1 fragment at the junction and another inter-chromosomal translocation of chr12 and chr3 with a 2013bp long L1 fragment - where both SVs shared the same breakpoint on chr12 (**Figure 3b**). Mapping the L1 fragments to a canonical L1 element revealed adjacent L1 segments, suggesting a common somatic L1 source element on chr12 for these two L1 fragments embedded at the inter-chromosomal breakpoint junctions. Therefore, we likely observe the genomic scars of a somatic 5’-truncated L1 insertion, which subsequently mediated the occurrence of two inter-chromosomal translocations, both originating from the same somatic L1 locus on chr12. We subsequently detected a number of complex SVs with analogous structures in several patients, such as another inter-chromosomal translocation involving chr4 and chr5 with a 804bp L1 fragment at the breakpoint, an inverted duplication with L1 traces at the breakpoint and further SVs with L1 fragments of different lengths **(Supplementary Figure 10b**).

We next extended our L1 analyses to the full cohort, where we detected a broad variation in the number of somatic L1 insertions between patients (0 to 542; **Supplementary Table A**). To validate the L1 activity beyond the promoter methylation status, we stained for L1 open reading frame 1 protein (ORF1p) on matched tissue sections (**Figure 3e**). We observed a correlation between ORF1p protein expression (signal intensity) and the number of L1 insertions detected using ONT (p = 0.084, r = 0.754, Spearman correlation; **Supplementary Figure 10h**), in agreement with occasionally extensive somatic L1 activity in bladder cancer samples.

As we identified somatic L1 insertions at SV breakpoints in B42, we wanted to explore whether L1s were significantly associated with markers of genomic instability across the cohort. We found a linear relationship between the count of non-insertion structural variants and somatic L1 insertions across tumors (p = 0.009, r = 0.53, Spearman correlation; **Figure 3f**). Notably, this relationship was also evident in the PCAWG cohort, where several tumor types showed strong associations between L1 insertions and structural variants. Head-SCC exhibited the strongest correlation (r = 0.75, p = 2.8e-11, Spearman correlation; **Figure 3g**), followed by other tumor types such as ProsAdeno-CA (r = 0.42, p = 1.2e-09, Spearman correlation; **Figure 3g, Supplementary Figure 10g**). Furthermore, tumors with high amplicon counts (≥ 20 amplicon regions) showed frequent somatic L1 insertions (**Figure 3h**, p = 0.0019, Rank-Biserial correlation (effect size) = 0.947, one-sided Mann-Whitney U-test), a pattern that extended to ecDNA presence (p = 0.0495, Rank-Biserial correlation (effect size) = 0.446, one-sided Mann-Whitney U-test; **Supplementary Figure 10f**). This pattern was exemplified by patient B123, which harbored high numbers of L1 insertions, amplicons, and multiple distinct ecDNA structures. Immunofluorescence analysis identified cells exhibiting both high *E2F3* (an oncogene present on ecDNA in B123) and L1 ORF1p expression (**Supplementary Figure 7g, Supplementary Figure 8**).

Thus, L1 insertions in our cohort were significantly associated with global genomic instability (i.e., complex SV rearrangements, high counts of SVs, amplicons and ecDNA presence). To further explore this relationship, we next performed a detailed spatial analysis of the most genomically unstable samples - those with ecDNA.

### Detection of ecDNA in space identifies ecDNA-specific transcriptional patterns

We observed evidence for greater genomic instability in ecDNA-positive samples, including elevated CNV scores (p < 0.0001, Rank-Biserial correlation (effect size) = -0.6, two-sided Mann-Whitney U-test; **Supplementary Figure 6b**) and higher counts of genetic clones (p = 0.12, Rank-Biserial correlation (effect size) = -0.714, two-sided Mann-Whitney U-test; **Supplementary Figure 6c**). We first developed an approach to detect ecDNAs within the tissue architecture from spatial transcriptomics data (see **Methods**). Using ONT data as a ground truth for ecDNA structures (initially for patient B4, due to the high tumor content for this patient), we showed a large range in the estimated copy-number of the ecDNA amplicons between and within the spatial clusters (**Figure 4a**). We established patient- and ecDNA-specific signatures to detect ecDNAs from Visium data (**Figure 4a**, **Supplementary Figures 11-14**, see **Methods**). ecDNA-high tissue regions corresponded to a subset of tumor tissue spatial areas (defined by SpatialDE2) (**Figure 4c-e**). Applying the ecDNA detection workflow to tumors B4 (**Figure 4, Supplementary Figures 11-12**), B42 (**Supplementary Figure 13**), and B123 (**Supplementary Figure 14**), which all harbored ecDNAs, revealed that ecDNA-specific signatures can be detected across patients. Hence, beyond bladder cancer, this new approach will help understanding ecDNA biology and studying ecDNAs in their spatial context, by linking ecDNA levels across subsets of tumor and non-tumor cells as well as across distinct spatial clusters.

**Figure 4.**
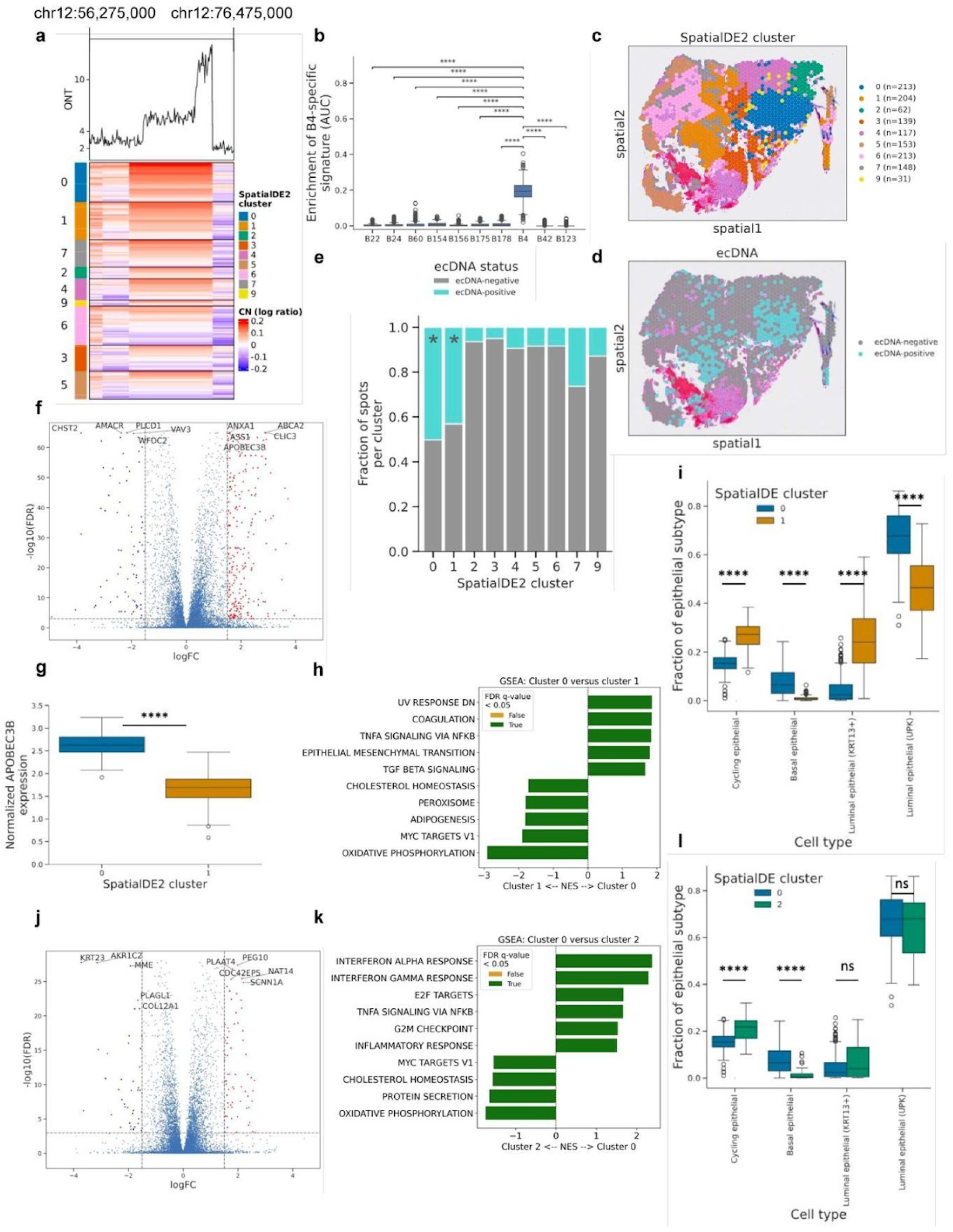
Detection of ecDNA in space identifies distinct tissue regions with different dosage of *APOBEC3B*. **a)** ecDNA detection using CNAs inferred from ONT (top) and CNAs inferred from Visium (tumor B4). Genomic region chr12:56,275,000-76,475,000 is shown. The annotation on the left represent transcriptionally similar regions identified by SpatialDE2, as shown in **c**. **b)** AUC scores (enrichment) of the gene set consisting of genes located on ecDNA in tumor B4, computed for each aneuploid spot in all 10 Visium samples. A non-parametric Kruskal–Wallis test was used for statistical testing, followed by Dunn’s test with Benjamini-Hochberg correction for multiple testing. Number of aneuploid spots per tumor: B22, n=504; B24, n=1171; B60, n=505; B154, n=2525; B156, n=1826; B175, n=1102; B178, n=1256; B4, n=1284; B42, n=1037; B123, n=1507. **c)** Spatial clusters detected by SpatialDE2, representing transcriptionally similar regions (number of spots per cluster are indicated). **d)** Predicted ecDNA status based on CopyKAT-inferred copy-number state of the ecDNA region and enrichment of the B4 ecDNA-specific gene set. **e)** Fraction of ecDNA-positive spots per SpatialDE2 cluster. Stars annotate the clusters significantly enriched with ecDNA-positive spots (one-sided Fisher’s exact test with Benjamini-Hochberg correction for multiple testing). Odds ratios for the main clusters: cluster 0 = 4.84; cluster 1 = 3.26; cluster 2 = 0.22. **f)** Volcano plot showing differentially expressed genes between cluster 0 and cluster 1 from **c**. **g)** Comparison of the normalized expression of *APOBEC3B* between ecDNA-enriched spatial clusters 0 and 1. Two-sided Mann-Whitney U-test was used for statistical testing. Rank-biserial correlation (effect size) = 0.989. **h)** GSEA on hallmark gene sets based on DEG analysis from **f**. **i)** Comparison of the abundance of epithelial cell subtypes between ecDNA-enriched clusters 0 and 1. Two-sided Mann-Whitney U-test was used for statistical testing with Benjamini-Hochberg correction for multiple testing. **j)** Volcano plot showing differentially expressed genes between cluster 0 and cluster 2. **k)** GSEA on hallmark gene sets based on DEG analysis from **j**. **l)** Comparison of the abundance of epithelial cell subtypes between cluster 0 and cluster 2. Two-sided Mann-Whitney U-test was used for statistical testing with Benjamini-Hochberg correction for multiple testing. All boxplots in Figure 4 show the median, interquartile range (IQR), and whiskers extending to 1.5× IQR.

In our cohort, applying this method for detailed spatial analysis of ecDNA in B4 revealed notable patterns to potentially explain the link between L1 insertions and ecDNA. Between the spatial areas with the highest enrichment of ecDNAs (clusters 0 and 1 for patient B4, respectively), we detected significant differences in the fractions of cell types and in the respective transcriptomes, with *APOBEC3B* as one of the most differentially expressed genes between these two ecDNA-high clusters (**Figure 4f-i, Supplementary Figure 6e** and **12**). GSEA analysis on hallmark gene sets identified TGF-alpha signaling, epithelial mesenchymal transition, and TGF-beta signaling as upregulated in cluster 0 (APOBEC3B-enriched) and oxidative phosphorylation and MYC targets as upregulated in cluster 1 (APOBEC3B-relatively reduced), respectively (**Figure 4h**). Importantly, APOBEC3B has previously been reported to be activated in the presence of L1 insertions to serve as an inhibitor of L1 retrotransposition^20,38^, and its function as an antiviral cytidine deaminase suggests a viral mimicry response mechanism. We hypothesize that enriched L1 retrotransposition activity within the APOBEC3B-enriched ecDNA-positive cluster 0 cells versus the APOBEC3B-relatively reduced ecDNA-positive cluster 1 cells might explain the differential expression of TNF-alpha and TGF-beta signaling.

More broadly, we also identified marked differences in cell type composition and biological processes when comparing ecDNA-positive with ecDNA-negative clusters (**Figure 4j-l**). Here, we again saw the upregulation of immune pathways including interferon alpha and interferon gamma response in the ecDNA-positive cluster 0 versus ecDNA-negative cluster 2. Importantly, ecDNAs have been associated with a dampened immune response, suggesting an alternative mechanism responsible for activating an immune response. We hypothesize that enriched L1 retrotransposition activity within the ecDNA-positive cluster 0 cells versus the ecDNA-negative cluster 2 cells could potentially contribute to this re-activated immune response.

Thus, from our spatial data we detected significant genomic instability via ecDNA linked to immune response, which was further associated with APOBEC3B-activity. To explore whether this result could potentially be mediated by L1s, we next sought to temporally relate L1s with genomic instability, and explore a viral mimicry response in our L1-high samples.

### Early L1 insertions leads to downstream genomic instability and viral mimicry response

To get insights into the temporal genomic evolution of the tumors, we first dissected clonal and subclonal events (**Methods**). Somatic L1 insertions were mostly clonal in this cohort (**Figure 5a**), as were the clock-like mutational signatures (**Figure 5b**). In contrast, signatures of APOBEC (SBS13, SBS2) and DNA repair (SBS3, SBS26) were largely subclonal. We also detected tobacco-smoking-associated signature SBS4 as predominantly subclonal (**Figure 5b, Supplementary Figure 15**). Given the reported association between *TP53* mutations and somatic L1 insertions^39,40^, including via an enhanced viral mimicry response pathway^41^, we also sought to explore possible relative timing between the events. As expected, *TP53*-mutant tumors showed significantly higher numbers of L1 insertions as compared to *TP53*-wild-type tumors (p < 2e-16, Rank-Biserial correlation (effect size) = 0.119, one-sided Mann-Whitney U-test, **Figure 5c)**. Of the 9 *TP53* mutation events identified, 7/9 were clonal (∼78%, **Supplementary Table H**). These results indicate *TP53* mutation and L1 insertion as an early, often clonal event, followed by other mutational processes including DNA repair and APOBEC-mediated editing, and a potentially enhanced viral mimicry response.

**Figure 5.**
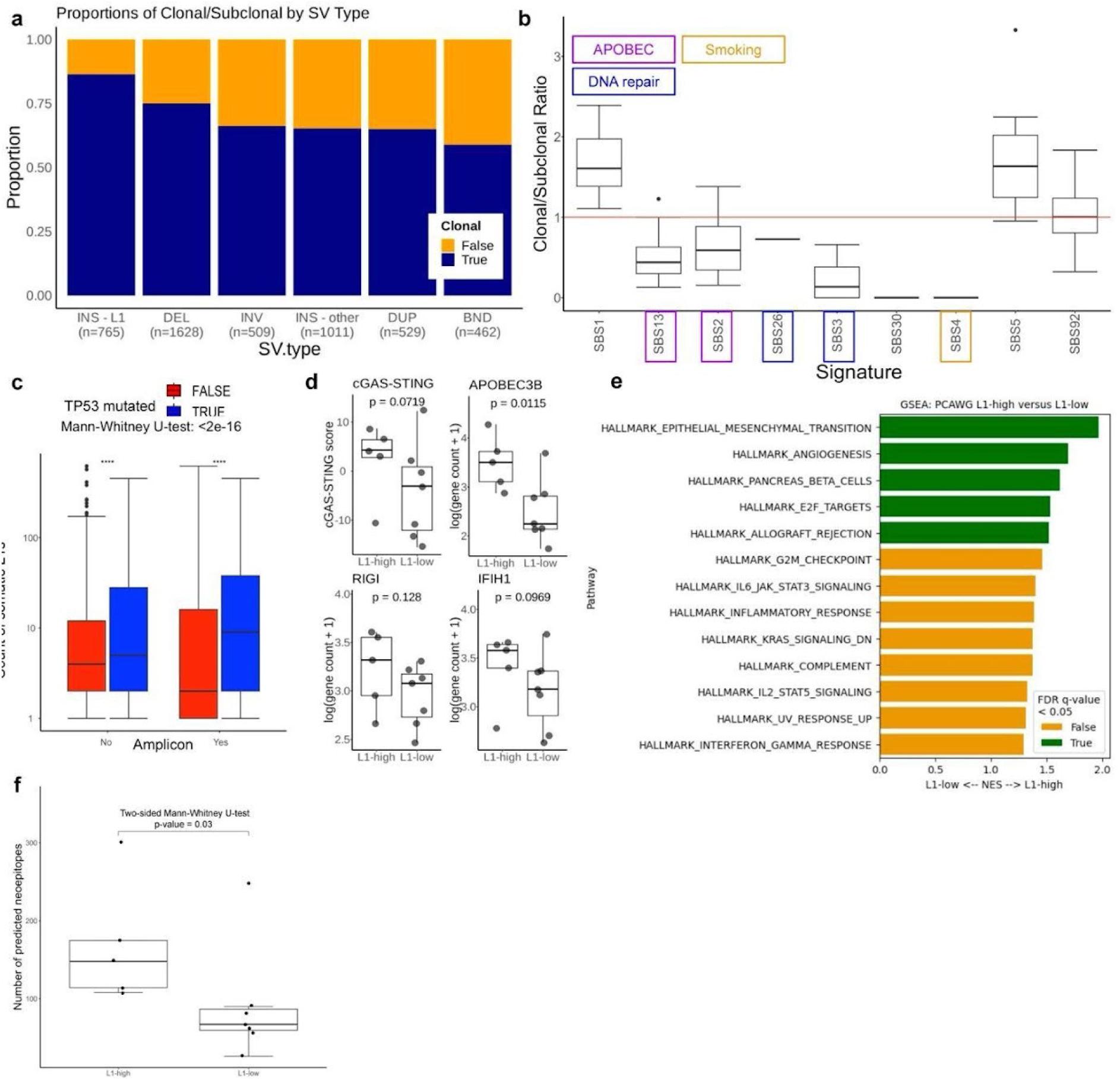
L1 insertions are clonal and correlated with mutational and DNA repair processes. **a)** Proportion of clonal structural variations (VAF > 0.3) across the full cohort, separated by SV type (ALU and SVA insertion types are excluded). **b)** Ratio of clonal to subclonal mutations grouped by mutational signature. Signatures with a ratio below the red line (h = 1) suggest late (subclonal) mutational events, while those above the line indicate early (clonal) events. **c)** Log-scaled count of L1 insertions in PCAWG dataset, split by presence of amplicons (Yes = 180098, No = 1166922) and TP53 mutations (True = 291405, False = 1055615). **d)** Expression of viral mimicry response pathway genes in L1-high versus L1-low samples, defined by insertion count thresholds (L1-high = insertion count > 25: n = 5, L1-low = insertion count < 3: n = 7). Significance assessed with one-sided Mann-Whitney U-test; see text for Rank-Biserial coefficients (effect sizes). **e)** Gene set enrichment analysis (GSEA) on hallmark gene sets of L1-high versus L1-low samples, defined by L1 insertion count percentiles (L1-high = top 10% across cohort: n = 3, L1-low = bottom 10% across cohort: n = 2). Sets with nominal p-value < 0.05 are displayed on the plot. Significance assessed by FDR q-value < 0.05 (green). **f)** Predicted neoepitope counts from single nucleotide variants and InDels from tumor samples with available RNA-seq data classified as L1-high (insertion count > 25, n = 5) and L1-low (insertion count < 3, n = 7). Statistical significance was evaluated using a two-sided Mann-Whitney U-test. All boxplots in Figure 5 show the median, interquartile range (IQR), and whiskers extending to 1.5× IQR.

To further explore the viral mimicry reaction suggested by our spatial transcriptome analyses, we next compared the expression of viral infection response pathway genes between our L1-high (L1 insertion count > 25, n = 5 with RNAseq data; B42, B123, B134, B4, B5) and L1-low samples (L1 insertion count < 3, n = 7 with RNAseq data; B24, B154, B178, B39, B60, B156, B74) (**Figure 5d**). The cGAS-STING score, a marker for viral infection response to cytosolic dsDNA, was increased in L1-high patients, although the difference did not achieve statistical significance with the available cohort size (p = 0.0719, Rank-Biserial correlation (effect size) = 0.543, one-sided Mann-Whitney U-test). *APOBEC3B* expression was significantly increased in the L1-high patients (p = 0.0115, Rank-Biserial correlation (effect size) = 0.829, one-sided Mann-Whitney U-test), supporting our previous spatial results. *RIG-I* and MDA5 (*IFIH1*) expression—markers of the viral response to cytosolic dsRNA—were both increased in L1-high patients; even though the differences were not statistically significant at this sample size (p = 0.128, Rank-Biserial correlation (effect size) = 0.429, and p = 0.0969, Rank-Biserial correlation (effect size) = 0.486, respectively; one-sided Mann-Whitney U-test). To support these findings, we ran GSEA on the L1-high versus L1-low samples using a stricter assignment of L1-high (top 10% of counts, n = 3; B42, B123, B134) and L1-low (bottom 10% of counts [both with zero L1 insertions], n = 2; B74, B156) samples (**Figure 5e**). We found significant L1-high enrichments of immune response pathways (allograft rejection, IL6 Jak/Stat signaling, inflammatory response, complement signaling, IL2 Stat5 signaling, Interferon gamma response) and cell growth/cell cycle progression gene sets (E2F targets, G2M checkpoint). In line with this, we also detected significantly more (p = 0.03, Two-sided Mann-Whitney U-test) predicted neoepitopes in L1-high as compared to L1-low patients (**Figure 5f**). Taken together, these results support an immune response to early, clonal L1 insertions.

## DISCUSSION

Our comprehensive multi-omic profiling of 49 bladder cancers revealed diverse patterns of genomic instability. Integrated analysis of copy-number and single-nucleotide variants revealed recurrent alterations in established bladder cancer driver genes, including deletions in *KDM6A* and *STAG2*, as well as high-frequency missense mutations in *NBPF10.* The long-read sequencing approach pursued in our study furthermore identified widespread structural variants and ecDNA, and particularly facilitated the discovery of somatic L1 insertion events across samples. These results were validated and further explored through spatial transcriptomics analysis, which identified ecDNA-specific transcriptional signatures within the tissue architecture and marked differences between ecDNA-positive tumor subclones, potentially caused by differential APOBEC activity. Timing analysis identified L1 insertions as an early, clonal event and our integrated analysis suggested genomic instability and viral mimicry response as downstream events.

In agreement with previous literature, clonal and subclonal mutation timing analyses showed L1 insertions to be mostly clonal^42^ and associated with *TP53* mutations^20^. Using long-read sequencing and de novo assembly, we identified active source elements and multi-jump L1 insertions, suggesting ongoing retrotransposition activity in tumors with hypomethylated L1 promoters. In contrast, the most confident L1-silent tumor in the cohort displayed methylated promoter regions at the same loci, reinforcing the role of epigenetic regulation. While L1 insertion length did not correlate with methylation status, we observed inter-patient variability in insertion burden, which aligned with ORF1p protein expression in tissues. These results confirmed the early and active nature of L1 insertions in our samples.

We identified multiple potential consequences of L1 retrotransposition in our patients. First, we showed a link between L1 insertions and the presence of genomic instability markers (including structural variants, amplicons, and ecDNA) in bladder cancer. Although direct evidence between L1 insertions and structural variant formation, especially ecDNA formation, is currently limited, several indirect associations suggest possible connections. First, L1 retrotransposition can cause genomic instability through insertions promoting double-stranded DNA breaks. This instability may facilitate formation of SVs and downstream ecDNAs; indeed, we saw evidence of several L1 fragments inserted at an SV breakpoint, and additional examples may potentially exist that are not yet detectable due to current sequencing technology and algorithmic limitations. While we did not observe evidence for L1 insertions at ecDNA breakpoints, we saw a significant association between L1 count and ecDNA presence. A causative relationship between L1 insertion and ecDNA formation is not implausible and should be explored in more detail; for example, a relationship between L1 insertions and the formation of breakage-fusion-bridge cycles, a common precursor to ecDNA, has been established in the PCAWG cohort^43^. In addition, ecDNA has been shown to function as mobile enhancers, interacting with chromosomal DNA to amplify gene expression^44^. These interactions could potentially include genes affected by L1 insertions, as previously described in a cancer context^37^, leading to coordinated regulation and amplification of specific genomic regions. Thus, it is important to continue exploring this question in more detail.

A second potential consequence of L1 retrotransposition that we found was the link between L1 insertions and APOBEC3B activity. APOBEC3B has been previously shown to be activated in response to L1 retrotransposition as part of an innate viral response^38^. In our dataset, *APOBEC3B* expression was significantly elevated in tumors with high L1 insertion counts. Based on this, we hypothesize that APOBEC3B may serve as a mechanistic link between L1s and ecDNAs. Supporting this hypothesis, a previous study identified co-occurrence of APOBEC3-associated kataegis and ecDNA - referred to as *kyklonas* - in 31% of tumors with ecDNA^25^. Notably, *kyklonas* were observed at a particularly high frequency in patients with Urothelial Bladder Carcinoma^45^. While this would suggest that *APOBEC3B* might be uniformly expressed across ecDNA positive spatial regions, our analysis revealed that *APOBEC* was among the most differentially expressed genes between regions positive for ecDNA. One possibility to explain differences in *APOBEC* expression may be mediated by differential L1 retrotransposition activity. In particular, because we identified APOBEC-related signatures as largely subclonal in our cohort, we propose a mechanism whereby early L1 insertions trigger an APOBEC3B response, which in turn support the editing and maintenance of ecDNA (the existence of which may potentially also be linked to L1 activity, as previously described).

Finally, we found that enriched *APOBEC* expression was just one component of a third potential consequence of L1 retrotransposition; an induced viral mimicry response. Overexpression of the *APOBEC3B* gene, alongside the cGAS-STING pathway^27^ and *RIG-I* and *MDA5* genes^26^, corroborates the viral mimicry response taking place in the L1-high samples. This is further supported by GSEA results, which show the significant overexpression of the interferon-alpha response pathway in L1-high samples. Hence, we link L1 retrotransposition activity to genomic instability and the viral mimicry response, highlighting the value of L1s as a potential biomarker for bladder cancers with complex tumor genomes. In addition, we hypothesize that early L1 insertions can trigger genomic instability events, including DSBs, and this can result in structural variants and potentially downstream ecDNA formation. We further hypothesize that the APOBEC3B-component of viral mimicry response to L1 retrotransposition may result in further maintenance and editing of ecDNA in these samples. These hypotheses should be further explored in future studies to clarify mechanistic relationships.

As an important side note, our analysis demonstrated substantial advantages of ONT sequencing for L1 discovery and epigenetic characterization, particularly in the context of complex SVs, where reads must span the SV junction and the embedded L1 fragment. However, we also frequently encountered mapping artifacts when L1s were inserted into the genome and long reads did not cover the entire insertion. This causes read aligners to map these fragments to alternative L1HS sequences present in the human genome leading to false positive SV signatures. Therefore, the development and extension of SV discovery methods is crucial in this context to safeguard long-read SV callers against such mapping artifacts and to identify the genuine retrotransposon-linked SV.

To the best of our knowledge, this is the first study of bladder cancer that integrates long-read DNA sequencing of tumors, short-read DNA sequencing of blood-based cfDNA, RNA sequencing, and spatial transcriptomics. This characterization of the genetic, epigenetic, transcriptomic, and spatial features of bladder cancer identified critical factors contributing to disease aggressiveness and suggested potential disease progression mechanisms, which will need further research to ultimately improve diagnosis, prognosis, and treatment for bladder cancer. In particular, we revealed evidence for previously suggested downstream effects of L1 retrotransposition. Our work highlights the value of L1 elements as a putative biomarker in bladder cancer and offers evidence for potential mechanisms of disease progression via L1 insertion-mediated genomic instability.

## ONLINE METHODS

### Material and data collection

All experiments in this study involving human tissue or data were conducted in accordance with the Declaration of Helsinki. Clinical data and tissue of all patients in the study were collected after receiving written informed consent from the respective patients or their legal representatives and after approval by the ethics committee of Heidelberg University. Participants did not receive compensation and were informed of this prior to enrolment in the study. Collection of blood and tumor material from patients with bladder carcinoma at diagnosis was done in collaboration with Dr. Mladen Stankovic and his team at the Salem hospital (Heidelberg).

### Blood collection

Blood was collected in cf-DNA/cf-RNA preservative tubes (Norgen Biotek) and mixed according to the manufacturer’s instructions. Blood tubes were collected from the Salem hospital and blood was processed immediately upon arrival at the laboratory. To separate plasma, blood tubes were centrifuged at 425xg for 20 minutes at room temperature. The upper plasma layer was carefully transferred by pipetting to clean Eppendorf tubes. Plasma aliquots were stored at -80°C until extraction of cfDNA. The rest of the blood pellet was saved and used for isolation of white blood cells (WBC) from which germline DNA was extracted.

### Extraction of cell-free DNA

Cell-free DNA extraction was done using Quick-cfDNA/cfRNA Serum & Plasma Kit (Cat#R1072, Zymo Research), following Parallel Purification protocol provided by the company. The standard input amount of plasma used for extraction was 2 ml. Lower amounts of plasma in exceptional cases of limited plasma material. The concentration of extracted cfDNA was measured with Qubit dsDNA High Sensitivity Assay Kit and length of cfDNA was determined by Bioanalyzer using Agilent High Sensitivity DNA Assay Kit.

### Library construction for cell-free DNA sequencing

Library construction was performed using KAPA HyperPrep Kit (Cat#7962363001, Roche), following manufacturer’s instructions. The standard cfDNA input amounts for library construction was 2.5-5.0 ng, depending on the quantity available. The order of steps followed from the manufacturer’s protocol was the following: End Repair and A-tailing, Adapter Ligation (Adapters used: KAPA Unique Dual-Indexed Adapter Kit; adapter concentration: 750 nM; end step incubation overnight at 16°C), Post-ligation Cleanup (step 3.14: 55 µl of Nuclease-free water used instead of the Elution buffer), Double-sided Size Selection (Appendix A1., using Ampure Xp beads Cat#A63881, instead of KAPA cleanup beads), Library Amplification, Post-amplification Cleanup. Final concentration of the libraries was checked with Qubit dsDNA High Sensitivity Assay Kit. Libraries were multiplexed and submitted for sequencing at the DKFZ NGS Core Facility. Whole-genome sequencing of cfDNA was done on Illumina NovaSeq 6000 S4 sequencing platform (100bp, paired-end).

### Cell-free DNA sequencing data analysis

FASTQ files were aligned to the GRCh38 reference genome with BWA-MEM (version 0.7.15)^46^. Duplicates were marked with picard tools (https://broadinstitute.github.io/picard/, version 2.25.1), which were removed in the next step in addition to low-quality reads, secondary alignments and supplementary alignments using SAMtools^47^. To enrich the tumor DNA, we only kept short fragments (90-150bp) as reported in previous studies^48^. Next, read counts were extracted using hmmcopy_utils (https://github.com/shahcompbio/hmmcopy_utils) for 500kb bin size. Finally, ichorCNA^49^ was used with default parameters to infer copy-number alterations and estimate the tumor fraction. One benign sample from our cohort was used as a reference. Only samples with the estimated tumor fraction above 0.03 were classified as positive.

### Tumor tissue collection

Tumor tissue (bladder carcinoma) was collected at diagnosis of each patient, during a procedure called Transurethral Resection of Bladder Tumor (TURBT). Removed tumor material was placed in a Falcon tube containing 5 ml of RPMI medium and transported on ice to the laboratory. The material was then frozen by dipping into pre-cooled 2-methylbutan for one minute, transferred into pre-cooled 1.8 mL Thermo Scientific CryoTube Vials and stored at -80°C until further cryosectioning.

### Cryosectioning and tumor content estimation

In order to confirm the presence of the tumor and estimate tumor content, we cryosectioned each tumor piece and performed Hematoxylin and Eosin staining, using standard procedures. If the tumor content was ≥ 70%, further cryosections were made for DNA extraction.

### Extraction of DNA for long-read sequencing

Extraction of DNA from cryosectioned tumor tissue was done using Gentra Puregene Tissue Kit (Cat#158063, Qiagen), following protocol for DNA purification from tissue using the Puregene Tissue Kit. In most cases, the amount of cryosectioned tumor material was between 5-30 mg tissue, therefore we followed the extraction protocol for low input amounts. For the elution step, a low EDTA buffer was used instead of the DNA hydration solution provided in the kit. The concentration of extracted tumor DNA was measured with Qubit dsDNA High Sensitivity Assay Kit and purity is analysed with NanoDrop. DNA was stored at -20°C until further use.

### Fragmentation of DNA

In order to obtain optimal DNA fragment length for long-read sequencing (10-30 kb), we performed the shearing of extracted high molecular weight DNA with the Megaruptor 3 system (EMBL, GeneCore Facility). The conditions used for shearing were the following: speed=30s, concentration=40ng/ul, vol=130ul. Shearing of DNA was followed by 1.0X v/v cleanup with SMRTbell beads (PacBio). Concentration of sheared DNA was measured with Qubit dsDNA High Sensitivity Assay Kit. Fragment length was analysed with the Agilent Femto Pulse system using Genomic DNA 165kb assay kit.

### Library construction for long-read DNA sequencing

Library construction was performed using Ligation Sequencing Kit V14 (SQK-LSK114, Oxford Nanopore), following the Library prep protocol from Oxford Nanopore called “Ligation sequencing DNA V14 (SQK-LSK114)”, further selected for PromethION device. Several steps of the protocol have been altered and all the alterations are listed below.

### DNA repair and end-prep

- Use of DNA Control Sample (DCS) was omitted and we used 48 ul of DNA (instead of 47 ul)
- Input DNA: 100-200 fmol
- Using a thermal cycler, we incubated at 20°C for 1 hour (instead of 5 min) and at 65°C for 10 min (instead of 5 min)
- Incubation on a Hula mixer was done for 30 min (instead of 5 min) at RT
- After resuspension of the pellet in 61 ul Nuclease-free water, we incubated for 10 min (instead of 2 min) at RT

### Adapter ligation and clean-up

- To enrich for DNA fragments of 3kb or longer, we used Long Fragment Buffer (LFB)
- Mix of DNA with Ligation buffer, Quick T4 DNA Ligase and Ligation Adapter was incubated for 20 min (instead of 10 min) at RT
- Incubation on a Hula mixer was done for 30 min (instead of 5 min) at RT
- After pellet resuspension in 25 ul Elution Buffer, the tube was incubated for 20 min (instead of 10 min) and at 37°C instead of RT

### Long-read tumor DNA sequencing

The input of the tumor DNA library used for loading on the flow cell was 20 fmol. We used R10.4.1 flow cells (FLO-PRO114M, Oxford Nanopore) and sequenced one tumor sample per flow cell. Sequencing was performed at the DKFZ Sequencing Open Lab on the PromethION (P24) device. Priming and loading of the flow cell was done according to Oxford Nanopore instructions. In order to reach optimal data output and coverage (30x), washing and re-loading of the flow cell was performed at least two times, approximately after 24h and 48h from the sequencing start. Total sequencing time was 96 hours.

### Germline DNA sequencing

DNA was extracted from white blood cells using the MonarchⓇ HMW DNA Extraction Kit for Cells & Blood (NEB, #T3050L) with the Standard Input protocol. Extracted DNA was sheared with the Megaruptor 3 system (EMBL, GeneCore Facility). The conditions used for shearing were the same as for tumor DNA: speed=30s, concentration=40ng/ul, vol=130µL. Shearing of DNA is followed by 1.0X v/v cleanup with SMRTbell beads (PacBio). Concentration of sheared DNA was measured with Qubit dsDNA High Sensitivity Assay Kit. Fragment length was analysed with the Agilent Femto Pulse system using Genomic DNA 165kb assay kit. Libraries for germline DNA bulk sequencing were done with the TruSeq Nano DNA LT LPK kit (order number 20015964) and the TruSeq DNA UD Index v2 (order number 20040870) both from Illumina. Samples were sequenced on a NovaSeq sequencing instrument in paired-end mode (2x150bp).

### Bulk RNA sequencing

RNA extraction was done with the Maxwell RSC Simply RNA Tissue Kit or with the RNeasy Kit (Qiagen) using the DNase treatment. The RNA quality was checked with a TapeStation. The library preparation was done at EMBL GeneCore. The RNA-seq libraries were prepared from total RNA with ribosomal RNA depletion using the NEBNext Ultra II Directional RNA Library Prep Kit for Illumina. Samples were multiplexed and sequenced using a NextSeq2000 sequencing instrument in paired-end mode. Depending on input RNA quality, samples were sequenced as 2x65bp (B39, B74 and B123) or 2x110bp (all other samples).

### Long-read basecalling and alignment

Basecalling of the Nanopore sequencing data was performed with Dorado version 0.6.0 (https://github.com/nanoporetech/dorado) using the most accurate PromethION basecalling model (sup) with modified base detection of 5-methylcytosine (5mC) and 5-hydroxymethylcytosine (5hmC) in CG contexts (sup,5mCG_5hmCG model). The base called data was then aligned to the human reference genome (GRCh38 and T2T) using minimap2^50^. T2T alignment was used to observe copy-number variations on chrY; GRCh38 alignment was used for remaining analysis.

### Long-read alignment quality control

Quality control of the long-read alignment files was done using NanoPack^51^, Alfred^52^ and verifyBamID^53^ to detect sample cross-contamination. The long read whole-genome coverage varied from 19x (B67) to 43x (B4) with an average coverage of 32x. The N50 read length varied from 11,565bp to 19,249bp with an average N50 read length of 15,655bp. The estimated sequencing error rate of the aligned data was in the range of 0.94% to 1.84% with an average estimated sequencing error rate of 1.17%.

### Single-nucleotide and small insertion and deletion calling

Clair3^54^ and ClairS (https://github.com/HKU-BAL/ClairS) were used to call germline and somatic single-nucleotide variants (SNVs) and small insertions and deletions (InDels), respectively. We used the default options of Clair3 and enabled phasing using WhatsHap^55^. We generated haplotagged BAM files using these phased SNVs and InDels to conduct allele-specific expression analyses with the nf-core RNA-Seq pipeline^56^ and allele-specific methylation analyses with modkit (https://github.com/nanoporetech/modkit).

### Structural variant calling

We used Delly^57^ and Severus^58^ to identify structural variants (SVs) from the long-read data. Because of our tumor-only long-read sequencing approach, we used a panel-of-normal (PoN) approach to distinguish germline and somatic SVs. For Severus and Delly, we used the 1000 Genomes ONT data generated by Schloissnig et al.^59^ for the generation of a PoN. Severus additionally requires phased SNVs, which we computed using Clair3^54^ (see above). We also used the default GRCh38 VNTR bed file provided by Severus for calling SVs. For delly, we used the default options in the long-read (lr) SV calling mode. To compute consensus SVs between Severus and Delly, we used Sansa’s compvcf subcommand (https://github.com/dellytools/sansa) to compare the output VCFs and identify matching somatic SVs between the two tools. We also used the matching short-read germline data for further filtering, where we computed SVs using Delly. To assign SVs to haplotypes, we split the BAM files by haplotype and re-run Delly on the haplotype-specific BAM files.

### Structural variant annotation

We used SVAN^59^ (https://github.com/REPBIO-LAB/SVAN) to annotate all consensus SV calls for variable number of tandem repeat (VNTR) variations, duplication variants and mobile element insertions (MEIs). We used the default options of SVAN for insertion and deletion annotation with the provided BED files for VNTR and mobile element annotation.

### Somatic retrotransposition analysis

To enhance specificity of our candidate somatic L1 callset, we called mobile element insertion in the matched short-read germline data using MELT^60^. We then compared all somatic L1 insertion with the germline L1 data using bedtools^61^ intersect on the insertion site with ±250bp to account for potential breakpoint ambiguities between the short- and long-read data. All candidate somatic L1 insertions overlapping a germline element were subsequently removed. To compare short- and long-read L1 calling, we used the matched tumor-normal B125 bladder cancer data. For the short-read data, we used MELT both on the tumor and matched control and compared the somatic L1s with the long-read based L1 calls for this sample. 6 out of 9 somatic L1s were directly confirmed by the short-read analysis. For the remaining 3, we used manual inspection in IGV^62^ to determine the somatic or germline L1 status. In one of the three events, we could not observe any abnormal mappings in the corresponding control, suggesting a somatic event. The other two candidate somatic L1s did show abnormal paired-ends in the control that were not called as an L1 by MELT but could be indicative of an insertion and hence, we labelled these events as unclear. An analogous comparison of the short-read based somatic L1 events called by MELT with long-read data showed an overlap of 6 out of 12 called SVs with only 2 long-read missed events classified as somatic after manual inspection in IGV^62^. One event was likely missed due to coverage (only 2 reads support) and the other event was a very short somatic insertion (81bp) that was called by Delly^57^ and Severus^58^ but not annotated as an L1 by SVAN.

### Retrotransposon-linked structural variants

Since Delly^57^ and Severus^58^ do not identify complex structural variants where somatic L1s can co-occur at the breakpoint of other SV types, we developed an algorithm to perform a targeted search for L1 sequence fragments at SV breakpoints. This new method, called BreakTracer (https://github.com/tobiasrausch/breaktracer) first identifies split-reads at SV breakpoints and then searches for L1 sequence fragments between these junctions to subsequently locally assemble all reads that support the same genomic breakpoints connected by an L1 insertion. We further filtered all breakpoints that co-occurred in multiple samples to prevent false positive retrotransposon-linked SV calls from mis-alignments or germline L1s that are not present in the reference genome.

### Copy-number variant calling

We used CNVkit^63^ and Delly’s cnv^57^ subcommand to call copy-number variants and generate read-depth profiles. For Delly, we used a window size of 25,000bp with the default GRCh38 mappability map downloaded from the Delly GitHub repository. For CNVkit, we selected the ‘wgs’ method and dropped low-coverage regions with the default GRCh38 reference file at 5k resolution.

### ecDNA identification

We used CoRAL^64^ to identify ecDNA structures from the long-read data with the copy-number profile computed by CNVkit^63^. We used the following parameter values: seed CN = 4.0, MBS = 2, CDA = 0.01. We further refined the structural predictions using orthogonal structural variant calls to select high-confidence ecDNA predictions.

### De novo assembly of source and somatic L1

We performed targeted de novo assembly of somatic L1 insertions using the Oxford Nanopore data for B42. Candidate insertion sites were extracted from the mobile-element annotated VCF file and extended by ±5 kb. Overlapping regions were merged, and reads mapping to these windows were extracted from the tumor alignment file using samtools^47^. Structural variants were called with Delly^57^ in long-read mode with the output of SV-supporting reads enabled, and insertions between 500–9,000 bp were selected. If a single variant was detected in a region, supporting reads were assembled using Shasta^65^. The resulting contigs were used to realign the original reads to the assembly, as well as a canonical L1 sequence from MELT^60^ using minimap2^50^ to confirm the L1 presence in the assembly. From these alignments, we inferred the L1 location in the contig using Alfred^52^ and its methylation status using modkit. We compared our source L1s against the published reference of repeat element L1s (UCSC Genome Browser hg38 RepeatMasker - LINE-1 track) and two additional source L1 lists^42,43^ to identify novel source L1s in our cohort. To identify “hot” L1 source regions, we merged overlapping L1 source elements and took the smallest start base pair and largest end base pair to define an L1 source region. These were checked against the previously cited reported L1 source element lists for overlap.

### Analysis of L1 regulation via methylation

To assess the methylation status of somatic L1 insertions in patient B42, we used modkit pileup to extract CpG methylation calls from Oxford Nanopore reads aligned to each assembled insertion. We then processed the resulting pileup files in R. After filtering for sites with coverage >5, we merged the methylation data with the corresponding L1 insertion metadata, including orientation and genomic coordinates. For full-length insertions (>6,000 bp), we defined the L1 promoter as the first 500 bp (strand-aware), and the L1 body as the remainder of the element. We then assigned each CpG to either promoter, body or 500bp flanking region and compared methylation levels between the regions using the two-sided Mann–Whitney U test. We next combined coordinate data from assembled L1 fragments (not full-length) and filtered out low-quality entries. L1 lengths were computed based on read alignment positions of a canonical L1 element (see above). Promoter regions were defined as the first 500 bp of each insertion, adjusted according to strand orientation. This annotated dataset provided the basis for integrating methylation profiles with insertion features.

### Circos Plot

L1 promoter and body regions were annotated using a curated Excel file and converted to BED format. Methylation data from nanopore sequencing pileup files were intersected with these regions using bedtools to extract CpG methylation calls. Average methylation fractions were calculated per region and linked to L1 insertion partner data. Copy number variation was assessed from tumor coverage files and integrated with methylation data for downstream analyses. The plot was created using the circlize R package.

### Visium spatial gene expression for FFPE tissue

Preparation of FFPE tumor tissue was done according to the 10X Tissue Preparation Guide. RNA quality assessment, section collection and placement, tissue adhesion test, probe hybridization and ligation, probe release and extension, library construction, and sequencing were done as described in our previous work^66^.

### Visium data processing and quality control

Raw visium FASTQ files were processed using the 10x Genomics SpaceRanger (version 2.0.1) with the GRCh38 reference genome. Downstream analyses were carried out using the Scanpy package (version 1.11.0)^67^. Spots with low number of counts (cutoff selected based on histograms) were removed. Genes detected in fewer than 10 spots were filtered out.

### Copy number inference from Visium spots

Copy number inference was performed using CopyKAT (version 1.1.0)^68^, with the following parameters: id.type="S", ngene.chr=5, win.size=25, KS.cut=0.1, distance="euclidean", output.seg="FALSE", plot.genes="TRUE", genome="hg20". Spots annotated as non-tumor by a pathologist were used as a reference for diploid cells via the norm.cell.names parameter. For samples where confident extraction of non-tumor spots was not feasible, the top 100 spots with the highest ESTIMATE score^69^ were used instead. CopyKAT-inferred copy number profiles were subsequently mapped to fixed 25 kb genomic windows for downstream analyses.

To define tumor subclones, we applied Leiden clustering to CNV profiles from all spots classified as aneuploid by CopyKAT. To determine the optimal Leiden resolution, we evaluated resolutions ranging from 0.1 to 0.4 and selected the one yielding the highest Calinski–Harabasz index. If the clustering solution consistently returned a single stable cluster across multiple resolutions, this was chosen as the final result.

Given that individual Visium spots may contain a mixture of tumor and non-tumor cells, the tumor-specific CNV signal can be diluted, potentially resulting in false subclone identification. To mitigate this, we computed a pairwise Pearson correlation coefficient between the pseudobulk CNV profiles of each subclone. Subclones with correlation coefficients above 0.95 were merged. Finally, for every spot we calculated CNV score defined as:

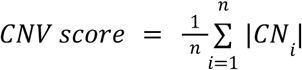

where *n* is the number of all genomic bins and *CN_i_* is the log ratio from CopyKAT at the *i*-th bin.

### Cell type deconvolution from Visium

Cell2location^70^ was used for cell type deconvolution. To compute the reference cell type signatures, the dataset by Gouin III et al. was used^71^ (GSE169379). As an alternative reference to validate the deconvolution, the dataset by Chen et al. was used^72^ (PRJNA662018). For the spatial mapping, we set hyperparameters to ‘N_cells_per_location = 20’ and ‘detection_alpha = 20’. Relative cell type abundance per spot was used for all the analyses.

### Detection of ecDNA in Visium spots

To predict the presence of extrachromosomal DNA (ecDNA) in Visium spots, we leveraged two key characteristics of ecDNA: (1) high-level focal genomic amplification, and (2) elevated expression of genes located on ecDNA.

First, to infer copy number profiles from Visium data, we used CopyKAT with the same parameters described previously, except for setting ‘win.size = 15’ to increase resolution. To validate focal amplifications, we used ecDNA structures reconstructed from ONT data to identify genomic regions known to be part of ecDNAs. We then examined whether these regions were also amplified in the CopyKAT CNV estimates. To classify gains in specific ecDNA-amplified regions for each spot, we fitted a Gaussian Mixture Model (GMM) with three components to the CopyKAT-derived copy number values. Spots assigned to the highest component were considered to show amplification in the ecDNA-associated region.

Next, we constructed sample-specific ecDNA expression signatures for all samples with both ecDNA and Visium data. These signatures consisted of genes mapped to the reconstructed ecDNAs. To retain only genes with elevated expression in the corresponding sample, we excluded any gene whose mean expression across aneuploid Visium spots was not highest in the sample being analyzed. We then computed signature enrichment scores using the ‘run_aucell’ function from the decoupler-py package (version 1.9.2)^73^. To account for differences in tumor purity and cellular composition, we regressed out the CNV score prior to analysis. Enrichment scores were binarized using the sample-specific median as a threshold. Spots were classified as ecDNA-positive if they showed both high signature enrichment and a gain in the ecDNA-amplified genomic region.

Finally, ecDNA status was propagated to transcriptionally similar spatial clusters identified by SpatialDE2^74^. Statistical significance was assessed using a one-sided Fisher’s exact test followed by Benjamini-Hochberg correction for multiple testing. Differential gene expression between clusters was performed in Scanpy with the function rank_genes_groups, using the two-sided Mann–Whitney U test, followed by gene set enrichment analysis (GSEA).

### Bulk RNA data analysis

For RNA-Seq quality control and gene quantification we used the nf-core RNA-Seq pipeline^56^. Reads were trimmed using TrimGalore (https://github.com/FelixKrueger/TrimGalore) and aligned using STAR^75^ version 2.7.11 to the GRCh38 reference genome. Salmon^76^ version 1.10 was used for computing gene counts and normalized transcripts per million (TPM) values. DESeq2^77^ was used for differential gene expression analyses of ecDNA positive samples compared to ecDNA negative samples and L1 positive samples compared to L1 negative samples. Pathway enrichment using cancer hallmark genes sets employed Gene Set Enrichment Analysis (GSEA). The top three L1-count samples were labeled as “L1-high”, and the bottom three L1-count samples were labeled as “L1-low”. The merged Salmon gene counts were then used to produce a ranked gene list for L1-high versus L1-low samples with DESeq2^77^ (v1.38.3). This ranked gene list was provided to GSEA (v4.4.0) for a GSEAPreranked run on Hallmark gene sets (h1.all.v2025.1.Hs.symbols.gmt).

For gene fusion identification, we used the nf-core RNA fusion pipeline^78^ and manually cross-checked predicted RNA fusions with the consensus somatic SV calls from the long-read DNA sequencing data. Gene fusion visualizations were generated using Arriba^79^.

### Identification of the driver genes in the patient cohort

We first performed a literature search to compile a list of established bladder cancer driver and marker genes. These genes were then screened in our patient cohort for copy number alterations (deletions, amplifications and loss of heterozygosity). All genes were visualized using the ComplexHeatmap R package.

### Clonality of structural variants and mutational signatures

Whole-genome sequencing data from clonal and subclonal single-nucleotide variants (SNVs) were analyzed to characterize mutational signatures in bladder cancer samples. SNV calls were parsed into sample-specific GenomicRanges objects using the GenomicRanges^80^ R package with genome built GRCh38 (hg38) as reference. Structural variant clonality was determined by calculating the variant allele frequency (VAF) for each SV, with a VAF>0.3 indicating a clonal rearrangement and VAF≤ 0.3 a subclonal rearrangement. Mutation type occurrences and 96-trinucleotide context mutation matrices were generated using the MutationalPatterns^81^ R package, using *fit_to_signatures_strict* function. Known COSMIC SBS mutational signatures (v3.2) were obtained and fitted to the mutation matrices to estimate signature contributions per sample via non-negative least squares regression implemented in MutationalPatterns.

Signature contributions were normalized to relative values within each tumor sample. Tumors with cosine similarity >0.95 between observed and reconstructed mutation profiles were selected for further analysis. Clonal-to-subclonal ratios of signature contributions were calculated per tumor, and aggregated ratios were visualized.

### Analysis of the L1 and TP53 status in PCAWG cohort

PCAWG samples were classified as “amplicon” or “no amplicon” based on CNV annotations (CN >6). L1 insertion counts were compared between amplicon and no-amplicon groups, stratified by *TP53* mutation status (mutant vs wild-type). Counts were log10-transformed for normalization.

### Fluorescence in situ hybridization

Two-colour interphase FISH was performed on human tissue sections using a rhodamine-labelled probe (centromere 6) and a FITC-labelled probe (*E2F3*; clone RP1-177P22). The probes were indirectly labelled via Nick translation. Pre-treatment of slides included use of Sodium thiocyanate (1 M NaSCN) at 80 °C. Digestion was performed with pepsin solution (1 mg/ml) for 30 min. Probes were applied on the slide, followed by a denaturation step for 10 min at 75 °C. Due to indirect probe labelling, signal detection was done on the next day, after washing slides with 50% formamide, followed by 0.5x SSC buffer. Samples showing sufficient FISH efficiency (>90% nuclei with signals) were evaluated.

### Ki67, LINE1-ORF1p and E2F3 stain and microscopy

For single staining, 4 μm thick formalin-fixed paraffin-embedded tissue sections were deparaffinized in xylene and subsequently hydrated. Antigen retrieval was carried out by the incubation of formalin fixed tissue in a 10 mM citrate buffer adjusted to pH 6.0 at 95–100 °C in a steam cooker for 40 minutes. The sections were blocked with 10% donkey serum diluted in 1x PBS. Anti-Line 1 ORFp1 antibody raised in mouse (Sigma, # MABC1152) at a dilution of 1:100 was used and kept for overnight incubation. Secondary donkey anti-mouse IgG Alexa Fluor 488 antibody (Life technologies, # A-21202) was used at a dilution of 1:500. Mounting of slides was performed with DAPI Fluoromount-G mounting medium (Southern Biotech, #0100-20).

For double staining of Ki67 and E2F3, 4 μm thick formalin-fixed paraffin-embedded tissue sections were deparaffinized in xylene and subsequently hydrated. Antigen retrieval was carried out by the incubation of formalin fixed tissue in a 10 mM citrate buffer adjusted to pH 6.0 at 95–100 °C in a steam cooker for 40 minutes. The sections were blocked with 10% donkey serum diluted in 1x PBS. Staining was done simultaneously for Anti-Ki67 raised in mouse (Agilent, M724029-2) diluted 1:100 and Anti-E2F3 raised in rabbit (Life technologies, PA5-106407) diluted 1:50. Antibody dilution was done with the Antibody diluent from Agilent (S3022). Secondary donkey anti-mouse IgG Alexa Fluor 488 antibody (Invitrogen, # A-21202) was used at a dilution of 1:500 and the secondary anti-rabbit Alexa Fluor Plus 594 (life technologies, A32754) at a dilution of 1:300. Vector® TrueVIEW® Autofluorescence Quenching Kit (#SP-8400) was used to quench autofluorescence from the tissue using Reagents A, B and C in equal volumes. Mounting of slides was performed with DAPI Fluoromount-G mounting medium (Southern Biotech, #0100-20).

For double staining of Line1 ORFp1 and E2F3, 4 μm thick formalin-fixed paraffin-embedded tissue sections were deparaffinized in xylene and subsequently hydrated. Antigen retrieval was carried out by the incubation of formalin fixed tissue in a 10 mM citrate buffer adjusted to pH 6.0 at 95–100 °C in a steam cooker for 40 minutes. The sections were blocked with 10% donkey serum diluted in 1x PBS. Staining was done simultaneously for Anti-Line 1ORFp1 raised in mouse (Sigma, # MABC1152) diluted 1:100 and Anti-E2F3 raised in rabbit (Life technologies, PA5-106407) diluted 1:50. Antibody dilution was done with the Antibody diluent from Agilent (S3022). Secondary donkey anti-mouse IgG Alexa Fluor 488 antibody (Invitrogen, # A-21202) was used at a dilution of 1:500 and the secondary anti-rabbit Alexa Fluor Plus 594 (life technologies, A32754) at a dilution of 1:300. Mounting of slides was performed with DAPI Fluoromount-G mounting medium (Southern Biotech, #0100-20).

### Confocal microscopy

Leica SP8 confocal microscope was used for the imaging of sections of FFPE tissue stained for LINE-1 ORF1p. Imaging was performed using the Leica LAS X software. Z-stacks were imaged with a step size of 0.5 μm. The start and end points of the Z-stack were set manually. Maximal intensity projections were made with Fiji with the Z projection function where the projection type was set to maximal intensity where all the stacks imaged were included in the projection.

### Imaging and scanning of human tissue sections

All H&E, Ki67, E2F3 and LINE-1 ORF1p stained sections were imaged using the Zeiss Axioscan 7 at the light microscopy facility at the DKFZ. Sections were imaged at 20x magnification with settings adapted from the fluorescence and brightfield profile for the respective fluorophores. Sample detection was performed by manual selection and focal points were assigned manually for each section.

### Identification of putative enhancer hijacking events using Pyjacker

Pyjacker is a computational method developed for the systematic prediction of enhancer hijacking events using various types of genomic data^31^. Pyjacker requires the following input files: a merged table of sample-specific structural variant breakpoints, corresponding gene expression values in TPM (Transcripts Per Million), and reference annotation files, including gene annotations and chromosome coordinates. Additionally, some optional inputs can be specified. In this study, the breakpoints table was generated from ONT (Oxford Nanopore Technologies) sequencing data and gene expression data were generated through RNA sequencing for the same samples. Furthermore, ChIP-seq datasets (ENCFF515VMS, ENCFF439WSM, ENCFF487EPL) were obtained from the ENCODE database^82^ to run the ROSE (Rank Ordering of Super-Enhancers) pipeline (https://bitbucket.org/young_computation/rose/src/master/) for identifying putative enhancers. Topologically Associated Domains (TADs) from Hi-C data were obtained from ENCODE (ENCFF682EJU) to complement the analysis. Pyjacker’s output is a table listing gene names, their genomic coordinates, and the corresponding false discovery rates (FDRs) for genes putatively affected by enhancer hijacking. To visualize the results, RStudio version 4.4.2 and figeno^31^ were used.

### Neoepitope prediction from somatic SNVs and InDels

Somatic single nucleotide variants (SNVs) and small insertions/deletions (InDels) called by Clair3 and ClairS were annotated with Ensembl VEP (v108, GRCh38) and annotated with expression data from matched RNA-seq (kallisto 0.51.1, Ensembl cDNA reference GRCh38, https://github.com/pachterlab/kallisto) using VAtools (https://github.com/griffithlab/VAtools). Only samples with available RNA-seq data were included in the analysis. HLA class I genotypes were determined by OptiType (https://github.com/FRED-2/OptiType) from short-read germline DNA. Neoepitope predictions for HLA class I were performed for peptides of 8-15 amino acids in length using pVACseq (pVACtools 5.3.0, https://github.com/griffithlab/pVACtools). Filtering steps included affinity thresholds (median IC50 < 500 nM or elution rank < 2%), variant quality (tumor DNA VAF > 25% and reads >10), and gene expression (>1 TPM). The output comprised filtered lists of predicted neoepitopes for each patient. Patients were grouped by L1 insertion burden, with >25 insertions labeled as high and <3 as low L1 insertion count. The number of predicted neoepitopes was compared between the L1 groups using a two-sided Mann-Whitney U test.

### Data visualization and statistics

For all data visualizations and statistics we used the following packages in R: dplyr, ggplot2, tibble, tidyr, stats, DESeq2, scales, DNAcopy, reshape2, grid, gridExtra, cowplot, ComplexHeatmap. We further used the following packages in python: gzip, pandas, matplotlib, numpy, glob, gseapy, scikit-learn, scipy, seaborn, pysam. Exact package versions can be found in the respective github pages listed in the Code Availability section.

## Data availability

Sequence data upload to the European Genome-phenome Archive (EGA) is in progress and will be available upon publication.

## Code availability

All open-source software code developed in this study is available via GitHub. The spatial and cfDNA analysis scripts are available at https://github.com/OtonicarJan/bladder-spatial. The long-read genomic data, expression and gene set enrichment analysis scripts are available at https://github.com/OsredekIvana/Bladder_cancer and https://github.com/sophiepribus/dkfz-embl-bladder. The method to screen long-read alignments for retrotransposon-linked SVs is available at https://github.com/tobiasrausch/breaktracer.

## ACKNOWLEDGEMENTS

We thank the team of Holger Sültmann at DKFZ and in particular Arlou Angeles and Florian Janke for advice on library preparation and data analysis. We thank Damir Krunic from the DKFZ light microscopy core facility for support with immunofluorescence signal quantification. We thank the DKFZ Sequencing facilities and EMBL GeneCore for excellent technical assistance, in particular Laura Villacorta for advice on ONT library preparation and Ferris Jung for RNA library preparation. We thank the DKFZ Single-cell open lab for support with the Visium experiments. We thank Frauke Devens and Michele Vousten for excellent technical assistance. We thank Petr Smirnov for discussions and help with data management. We thank Maike Hoesch, Jonas Bökelmann, Anne-Sophie Vieira Aleixo and Cecilia Martuzzi for support in the sample collection and experiments. AE received grants from the German Cancer Aid, the Mohr Foundation and the DFG.

## Author contributions

Conceptualization: S.J.P, I.O, J.O, M.S.L, J.O.K, T.R., and A.E. Clinical recruitment and support: M.St. Pathology analysis: K.B. Library preparation and sequencing: M.S.L., U.P., V.B., P.S., P.M. Bioinformatics and statistical analysis: S.J.P, I.O, J.O, and T.R. Analysis of the enhancer hijacking events: M.S. and S.M.-S. Analysis of the predicted neoantigens: A.K. Supervision of the neoantigen analysis: A.R. Advice on the cell-free DNA analyses: H.S. Advice on the enhancer hijacking analyses: C.P. Supervision: J.O.K, T.R., and A.E. Writing (original draft): S.J.P, I.O, J.O., T.R., and A.E. Writing (review and editing): all authors.

## Ethics declarations and competing interests

The authors declare no competing interests.

