## Supplemental Figures and Text for "Integrative spatial and multi-omic profiling in bladder cancer links L1 retrotransposition to extrachromosomal DNA, genomic instability, and viral mimicry response"

SUPPLEMENTARY FIGURES

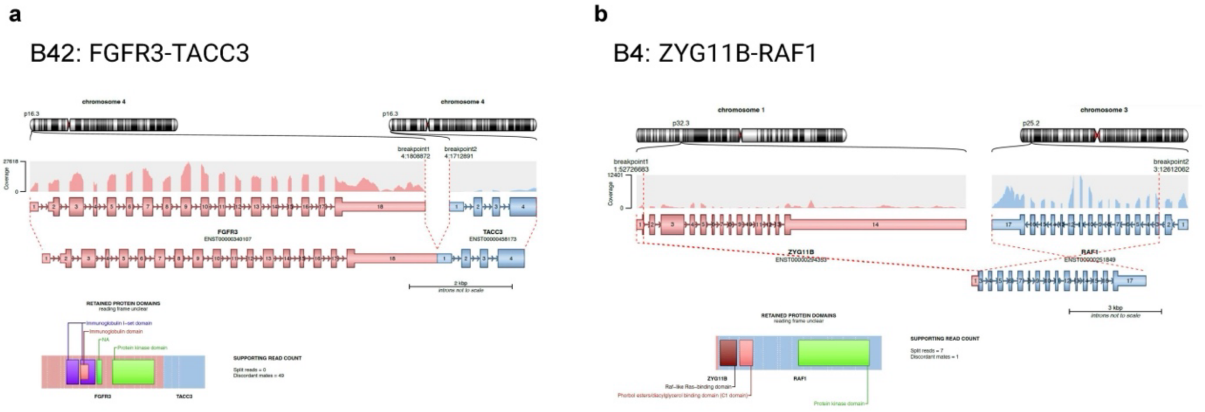


**Supplementary figure 1.** Arriba visualization of two highly expressed gene fusions FGFR3-TACC3 **a)** and ZYG11B-RAF1 **b)** in patients B42 and B5, respectively.

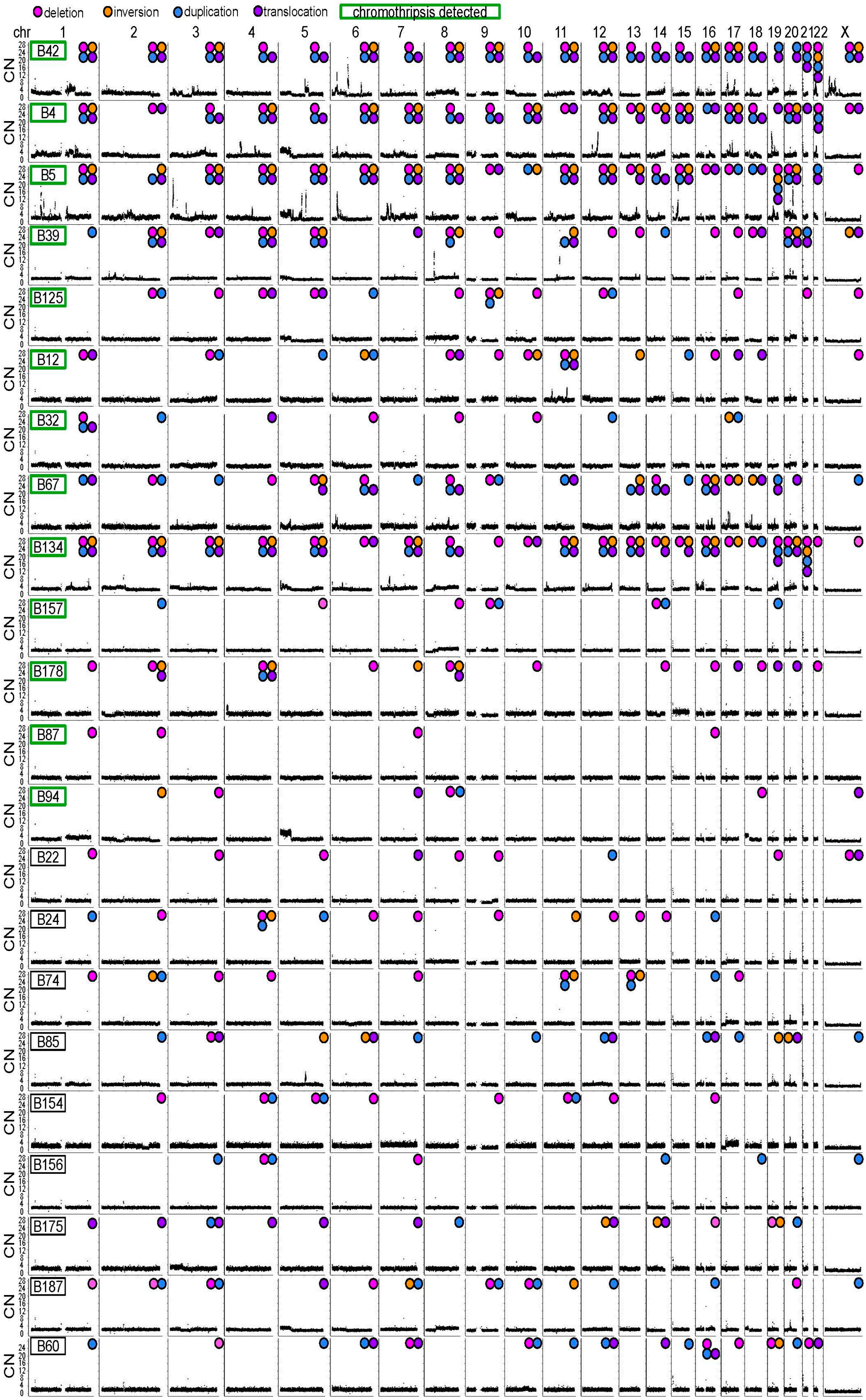
deletion
inversion
duplication
translocation
chromothripsis detected
chr 1 2 3 4 5 6 7 8 9 10 11 12 13 14 15 16 17 18 19 20 21 22 X
CN
B42
B4
B5
B39
B125
B12
B32
B67
B134
B157
B178
B87
B94
B22
B24
B74
B85
B154
B156
B175
B187
B60

**Supplementary figure 2.** Long-read WGS of bladder cancer samples reveals broad copy-number variation, sporadic high copy-number amplifications, and extensive structural variants. Chromosomes are labeled with the detected structural variation events. Samples scored positively for chromothripsis are labeled in green.

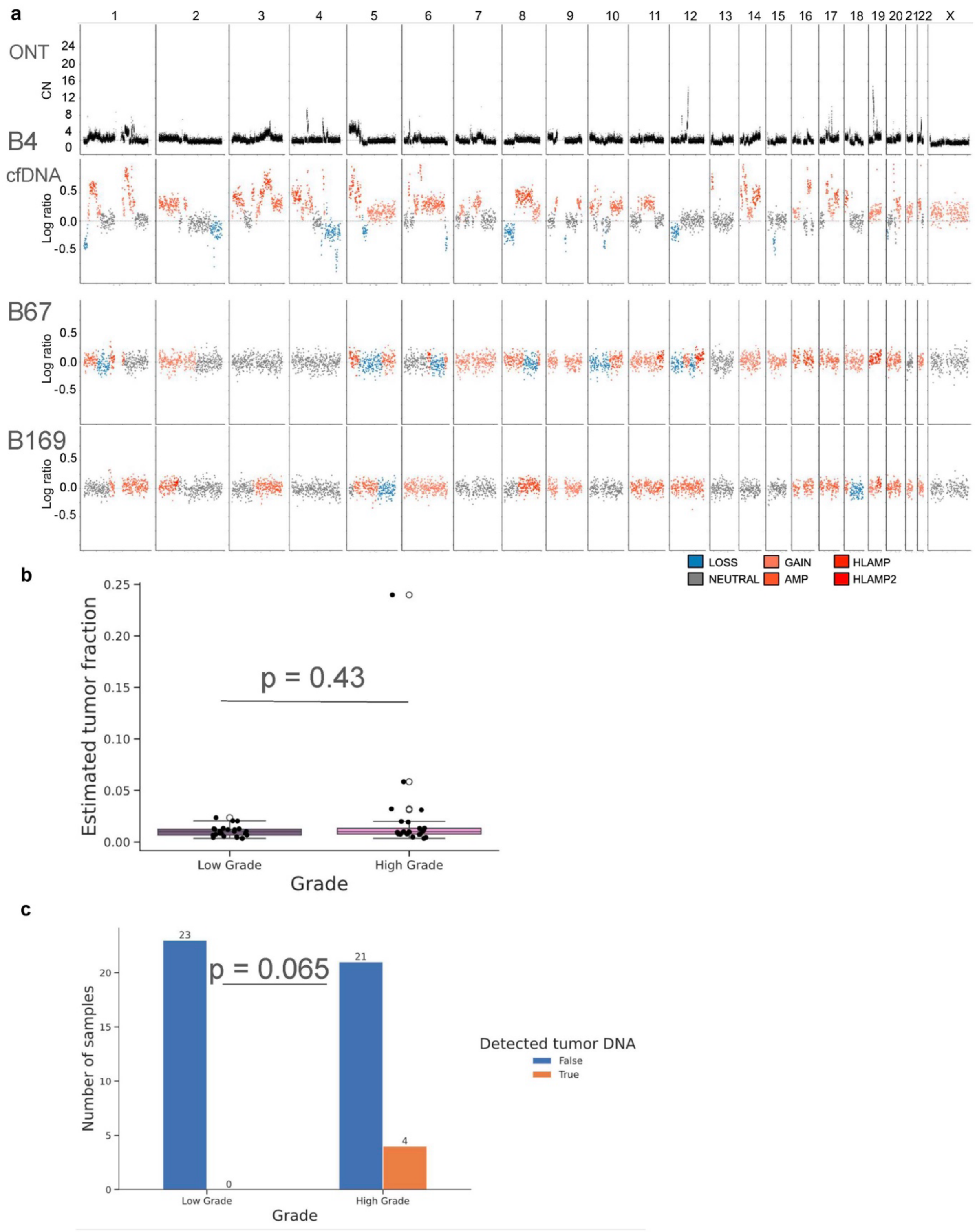


**Supplementary figure 3.** A minor fraction of cfDNA samples (4/25 high-grade tumors, 0/23 low-grade) show copy-number changes. **a)** Copy-number alterations for the sample B4 inferred from ONT or cfDNA, B67 (cfDNA only) and B169 (cfDNA only). LOSS: CN < 2, NEUTRAL: CN = 2, GAIN: CN = 3, AMP: CN = 4, HLAMP: CN = 5, HLAMP2: CN >= 6. **b)** Estimated tumor fraction from cfDNA for all 48 bladder cancer patients (the benign sample was not included). Two-sided Mann-Whitney-U test was used for statistical testing. Rank-biserial correlation (effect size) = -0.134. **c)** Number of samples with detected tumor DNA from liquid biopsy. A threshold of 3% estimated tumor fraction was used to classify detected tumor DNA. One-sided Fisher's exact test was used for statistical testing.

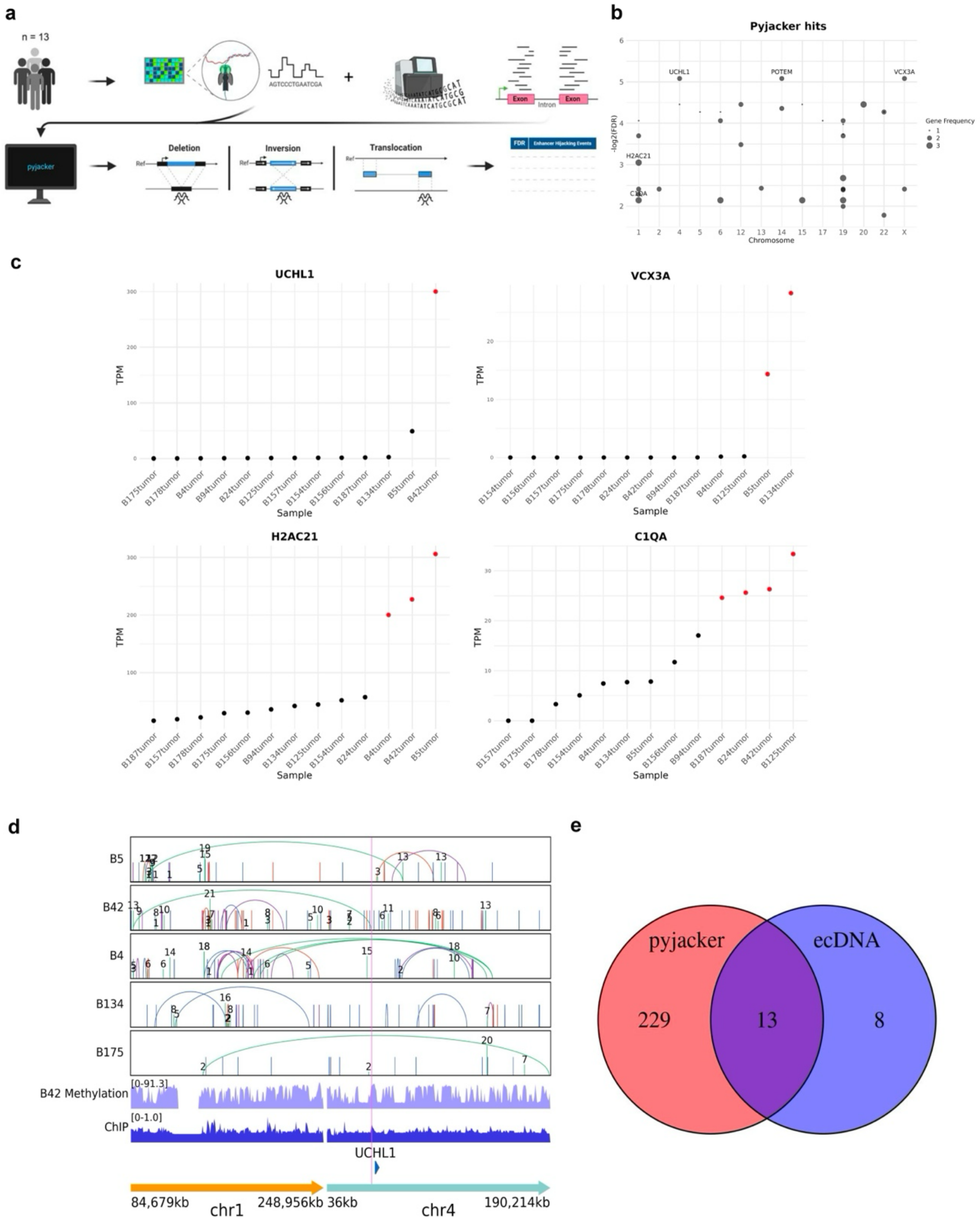


**Supplementary figure 4. a)** Workflow for the detection of putative enhancer hijacking events. **b**) Plot displaying the genes associated with the lowest FDR and most recurrent (defined by the number of samples in which the enhancer hijacking was detected) putative enhancer hijacking events as defined by pyjacker. **c**) Representation of gene expression in TPMs for four events with the lowest FDR and highest recurrence, with red points indicating the samples in which the gene is overexpressed (mean + 1 SD). **d**) Visualization of structural variations (SVs) in the UCHL1 gene across different samples. The SV involved in the putative enhancer hijacking event was a translocation in sample B42, shown here in green. Breakpoints are highlighted in pink. **e**) Venn diagram showing the overlap between genes overexpressed through enhancer hijacking as detected by pyjacker and through ecDNA, respectively.

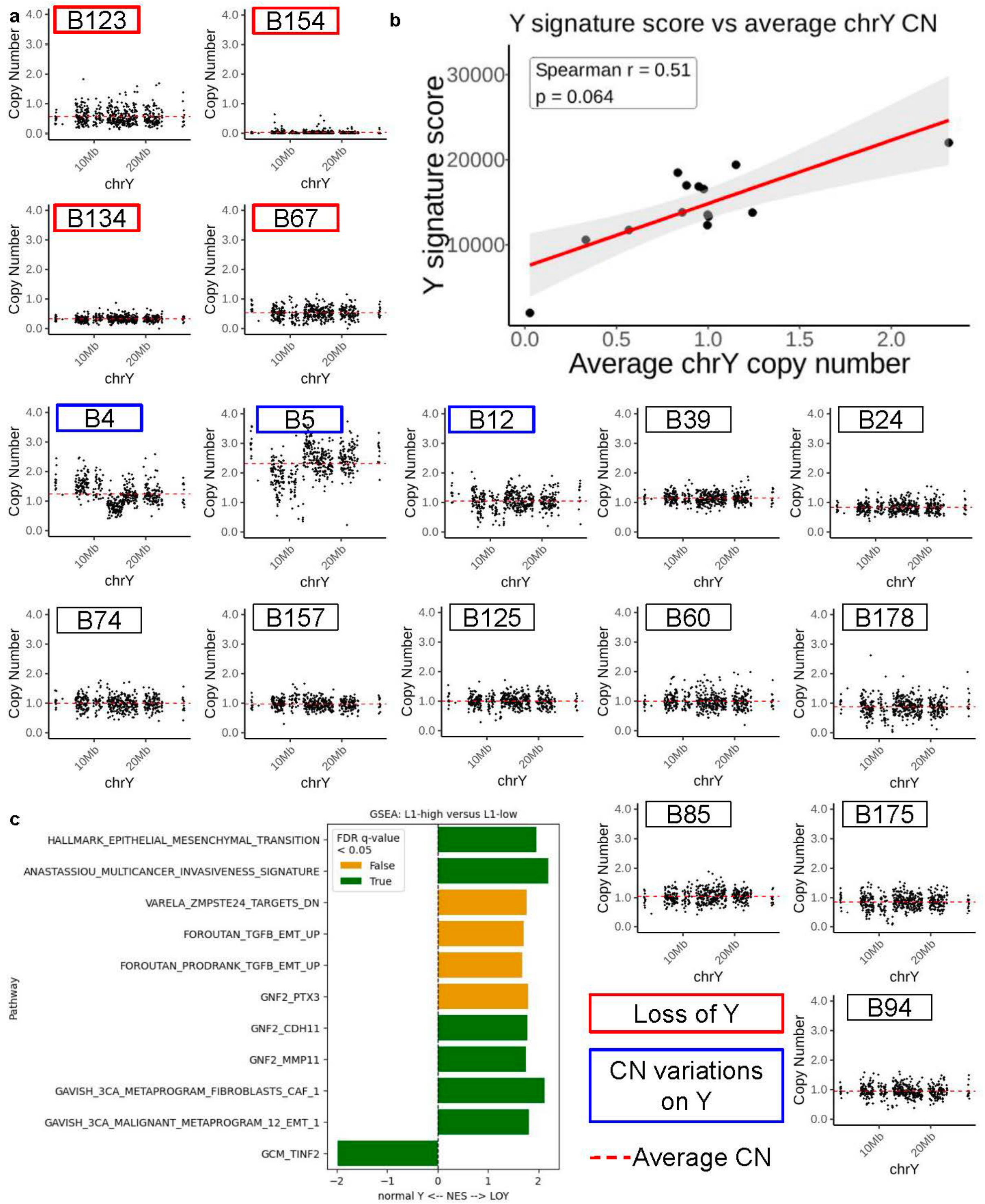


**Supplementary figure 5. a)** Copy-number plots across chromosome Y in male patients show heterogeneous loss of Y (n = 5), copy number variations (n = 3), gain of Y (n = 1), and normal Y expression (n = 10). **b)** ChrY signature score plotted against average chrY copy-number shows positive correlation between chrY copy number and chrY signature score. **c)** GSEA of multiple human gene sets shows significant enrichment of sets (nominal p-value < 0.05) involved in metastasis and invasion in LOY patients, versus tumor suppression in normal Y patients (note; GSEA run separately for each gene set collection [hallmark, c2-cgp, c4-cgn, c4-3ca], and specific gene sets from each selected for visualization). Significance assessed by FDR q-value < 0.05 (green).

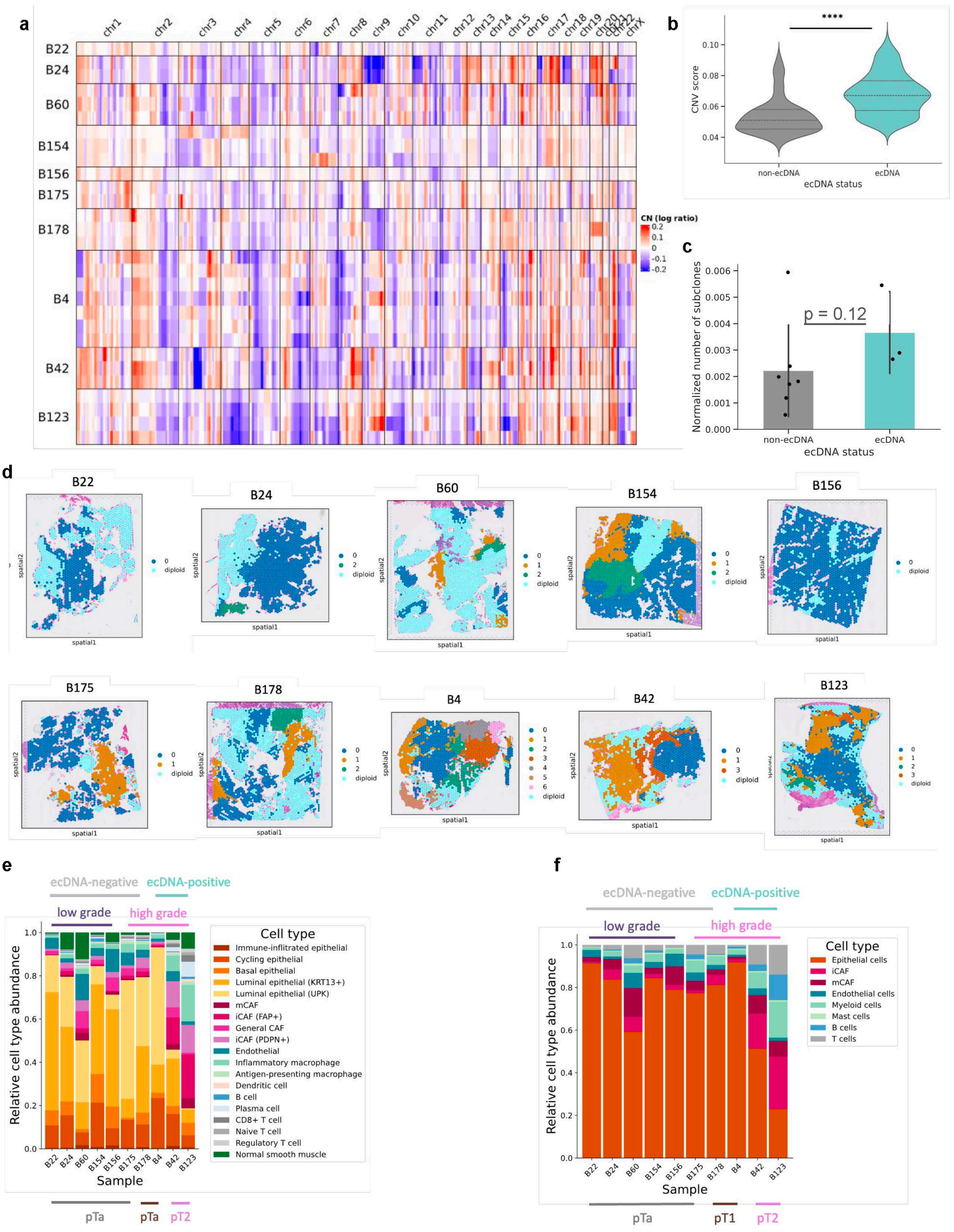


**Supplementary figure 6.** Copy-number inference and cell type deconvolution from 10 Visium samples. **a)** Heatmap showing pseudobulk CNA profiles of all the detected subclones for each tumor. **b)** Comparison in the CNV score for all aneuploid Visium spots between ecDNA-negative (n=8,889) and ecDNA-positive (n=3,828) tumors. Two-sided Mann-Whitney-U test was used for statistical testing. Rank-Biserial correlation (effect size) = -0.6. **c)** Comparison in the number of detected subclones from Visium between ecDNA-negative (n=7) and ecDNA-positive (n=3) tumors. Two-sided Mann-Whitney-U test was used for statistical testing. Rank-Biserial correlation (effect size) = -0.714. Height of the bars represent the mean, while error bars represent standard deviation. Each data point represents one tumor. **d)** Subclones identified by CopyKAT mapped to the Visium slides. **e)** Cell type composition of the Visium cohort, following deconvolution of cell types with cell2location, using a single-nuclei RNAseq reference from Gouin III *et al.,* or **f)** from single-cell RNAseq reference from Chen *et al.*

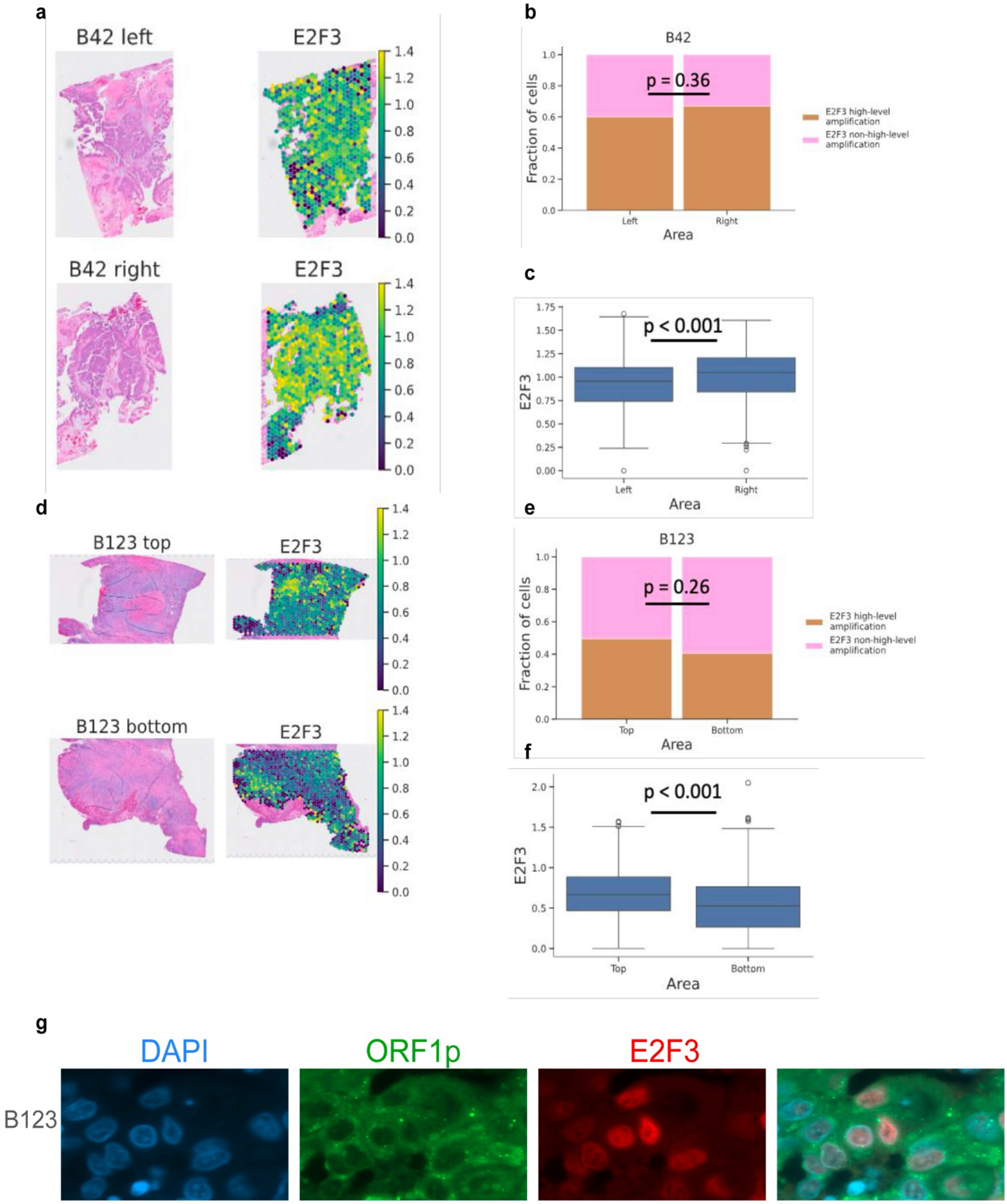


**Supplementary figure 7.** Quantification of high-level amplification of E2F3 by FISH validate Visium analysis results. **a)** Two regions of sample B42 used for FISH (left) with normalised E2F3 expression from Visium (right). **b)** Fraction of cells with E2F3 high-level amplification quantified by FISH for two distinct regions in sample B42. Chi-squared test was used for statistical testing. Number of counted cells per region: left, n=110; right, n=100. **c)** Normalised E2F3 expression from Visium for two distinct regions in sample B42. Two-sided Mann-Whitney-U test was used for statistical testing. Rank-Biserial correlation (effect size) = -0.21. Number of spots per region: left, n=737; right, n=687. **d)** Two regions of tumor B123 used for FISH (left) with normalised E2F3 expression from Visium (right). **e)** Fraction of cells with E2F3 high-level amplification quantified by FISH for two distinct regions in sample B123. Chi-squared test was used for statistical testing. Number of counted cells per region: top, n=101; bottom, n=101. **f)** Normalised E2F3 expression from Visium for two distinct region in sample B123. Two-sided Mann-Whitney-U test was used for statistical testing. Rank-Biserial correlation (effect size) = 0.234. Number of spots per region: top, n=1006; bottom, n=1276. **g)** Co-expression of E2F3 and LINE1 ORF1p in the same cells. Representative immunofluorescence stainings for the sample B123 are shown.

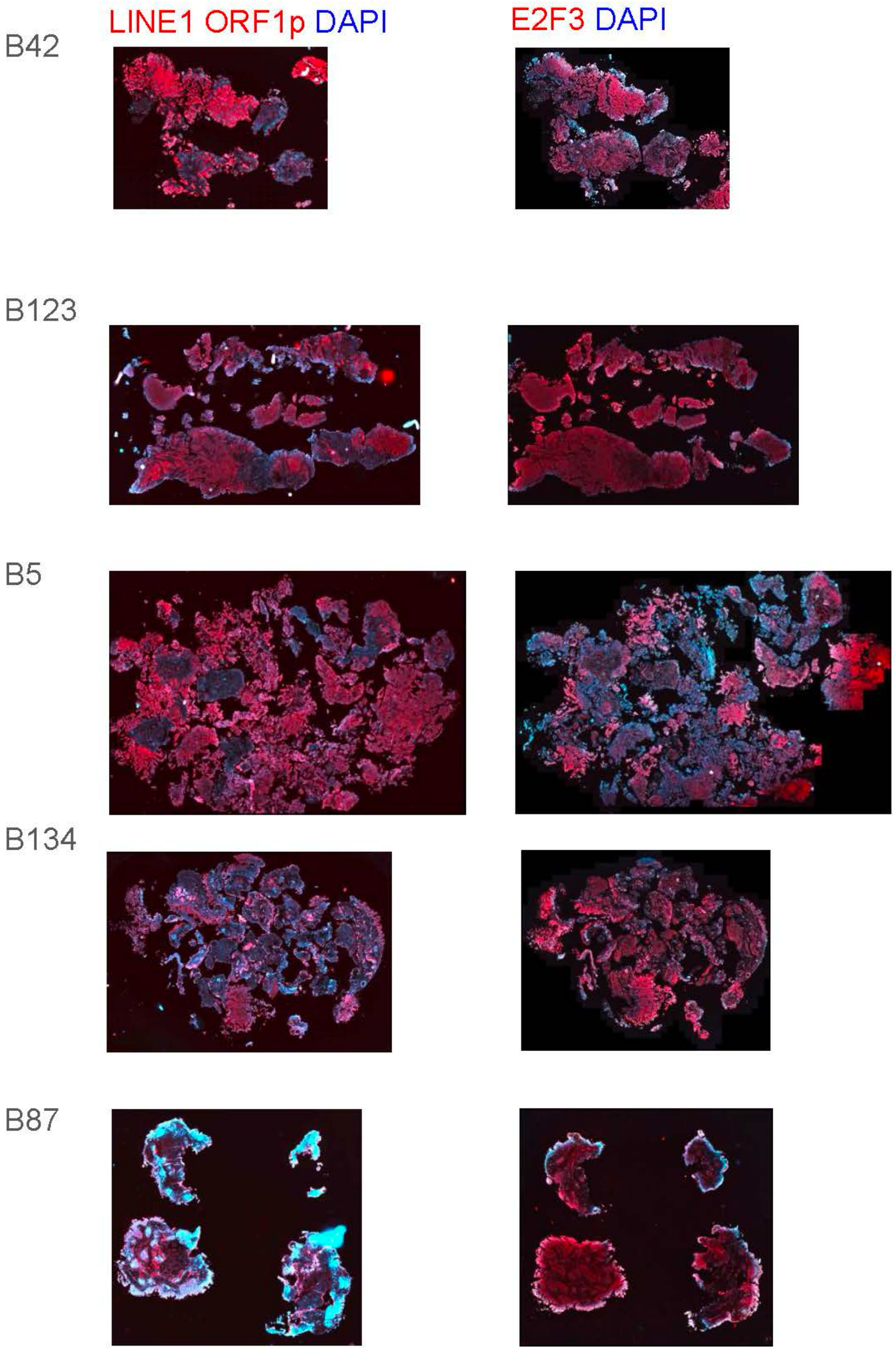


**Supplementary figure 8.** Expression of E2F3 (oncogene located on a ecDNA structure) and LINE1 ORF1p in the same tissue regions. Representative immunofluorescence stainings are shown.

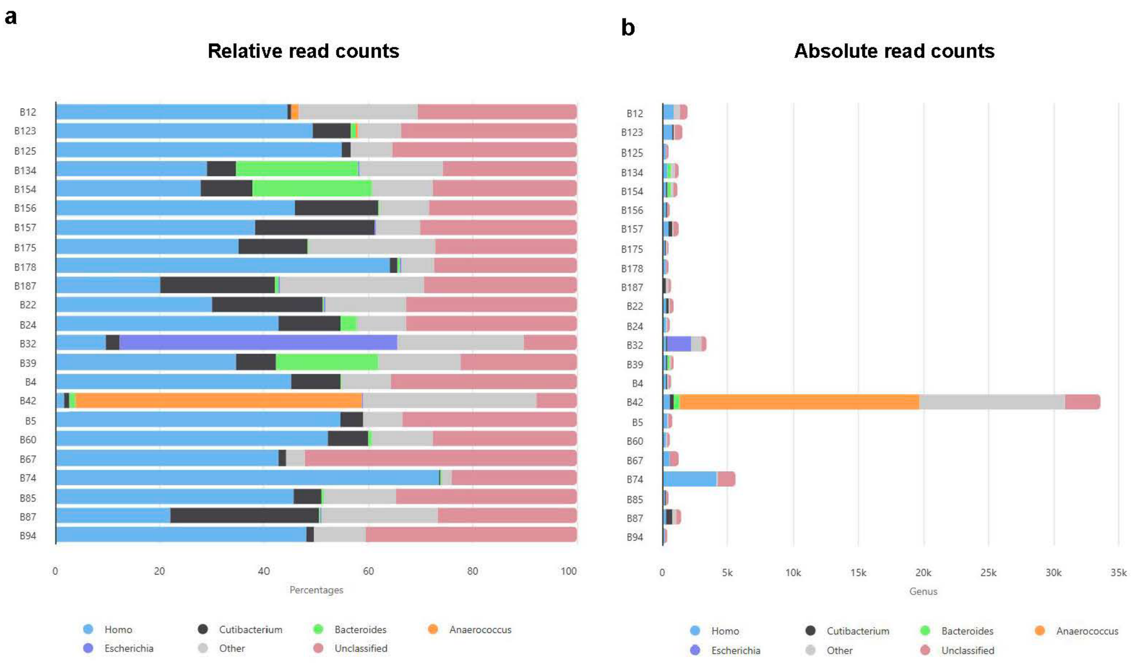


**Supplementary figure 9. a)** Percentages of unclassified and classified reads at the Genus taxonomy level. **b)** Absolute read counts. The analysis was performed on all unmapped long-reads from the tumor sample using kraken2.

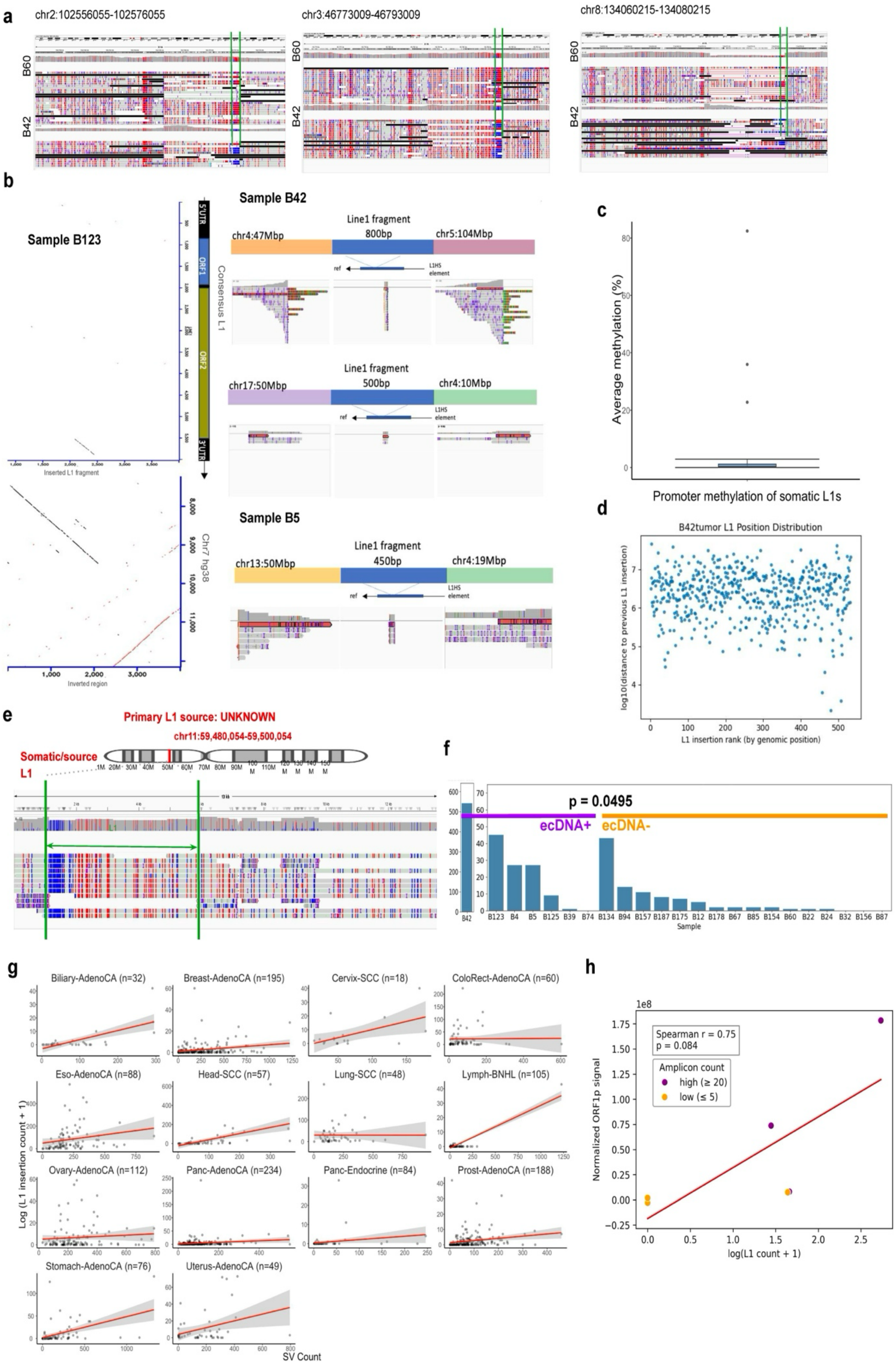

a
chr2:102556055-102576055
chr3:46773009-46793009
chr8:134060215-134080215
B60
B42
b
Sample B123
Consensus L1
5'UTR
ORF1
ORF2
3'UTR
Inserted L1 fragment
Chr7 hg38
Inverted region
Sample B42
chr4:47Mbp
Line1 fragment 800bp
chr5:104Mbp
chr17:50Mbp
Line1 fragment 500bp
chr4:10Mbp
Sample B5
chr13:50Mbp
Line1 fragment 450bp
chr4:19Mbp
ref
L1HS element
c
Average methylation (%)
Promoter methylation of somatic L1s
d
B42tumor L1 Position Distribution
log10(distance to previous L1 insertion)
L1 insertion rank (by genomic position)
e
Primary L1 source: UNKNOWN
chr11:59,480,054-59,500,054
Somatic/source L1
f
p = 0.0495
ecDNA+
ecDNA-
B42
B123 B4 B5 B125 B39 B74 B134 B94 B157 B187 B175 B12 B178 B67 B85 B154 B60 B22 B24 B32 B156 B87
Sample
g
Biliary-AdenoCA (n=32)
Breast-AdenoCA (n=195)
Cervix-SCC (n=18)
ColoRect-AdenoCA (n=60)
Eso-AdenoCA (n=88)
Head-SCC (n=57)
Lung-SCC (n=48)
Lymph-BNHL (n=105)
Ovary-AdenoCA (n=112)
Panc-AdenoCA (n=234)
Panc-Endocrine (n=84)
Prost-AdenoCA (n=188)
Stomach-AdenoCA (n=76)
Uterus-AdenoCA (n=49)
Log (L1 insertion count + 1)
SV Count
h
1e8
Spearman r = 0.75
p = 0.084
Amplicon count
high (≥ 20)
low (≤ 5)
Normalized ORF1p signal
log(L1 count + 1)

**Supplementary figure 10. a)** Comparative CpG methylation profiles of hg38-annotated LINE-1 loci in patient samples B60 (low L1 activity) and B42 (high L1 activity). Promoter regions are indicated with green bars. 20,000 bp genomic region in which L1 elements are found are indicated by the coordinates above each IGV plot. The 2-color 5mC IGV mode visualizes unmethylated bases in blue and 5-Methylcytosine (5mC) in red. **b)** Examples of a somatic L1 insertion embedded between somatic structural variant (SV) breakpoints in samples B123, B42 and B4. **c)** Average promoter methylation of the full-length somatic L1 elements in B42 (n=16). **d)** Rainfall plot of L1 insertion positions across the genome in B42 tumor. **e)** Second example of a multi-jump L1 event detected in the B42 tumor, visualized in IGV. A solo L1 insertion without any transduced sequence on chromosome 11 is visualized with unmethylated CpG sites shown in blue. **f)** Counts of L1 elements across tumors. ecDNA-positive tumors ($n = 7$) show a significant enrichment for L1 insertions versus ecDNA-negative tumors ($n = 16$) ($p = 0.0495$, Rank-biserial effect size = 0.446, one-sided Mann-Whitney U-test). **g)** Correlation between the number of L1 insertions and non-insertion SVs across PCAWG for tumors with more than 10 samples and at least one tumor with L1 count > 25. **h)** Correlation between the number of LINE-1 insertions and ORF1p protein expression (signal intensity). Each data point shows one tumor.

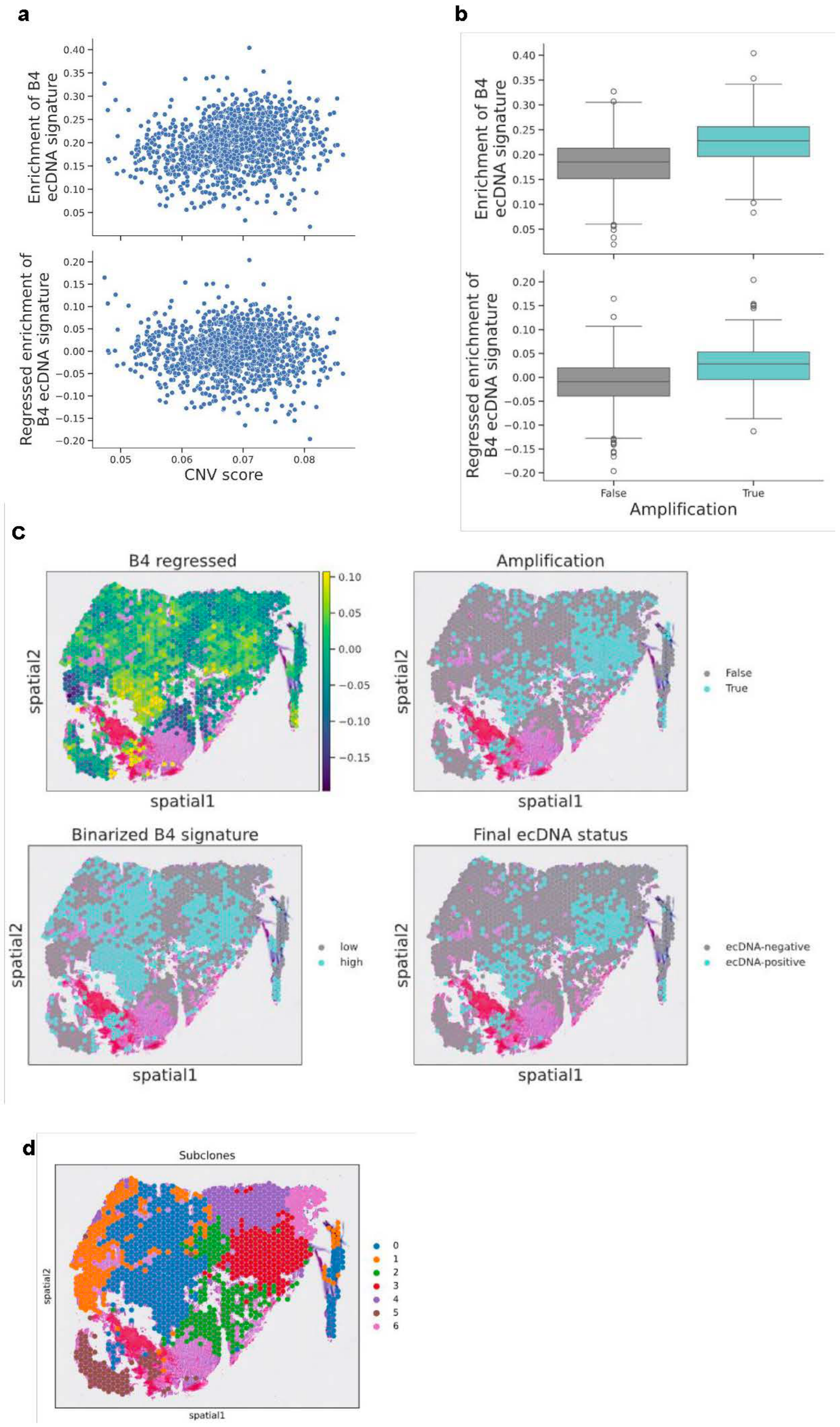


**Supplementary figure 11. Detection of ecDNA in Visium data for the sample B4. a)** Top, relationship between CNV score and AUC score of B4 ecDNA-specific signature. Bottom, relationship between CNV score and AUC score of B4 ecDNA-specific signature with CNV score regressed out per spot for the sample B4. **b)** Comparison in the enrichment of B4 ecDNA-specific signature between the spots without detected amplification at the genomic region with ecDNA and spots with amplification at the genomic region with ecDNA (top). The bottom boxplot shows the same, but with the enrichment of B4 ecDNA-specific signature that was regressed out by CNV score. **c)** Enrichment of the corrected B4 ecDNA-specific signature in space (top left), detection of amplification at the ecDNA genomic region per Visium spot (top right), corrected and binarized B4 ecDNA-signature (bottom left), the final ecDNA status predictions per spot based on consensus calls for the amplifications and high ecDNA signature enrichment (bottom right). **d)** Subclones derived from CopyKAT CNAs.

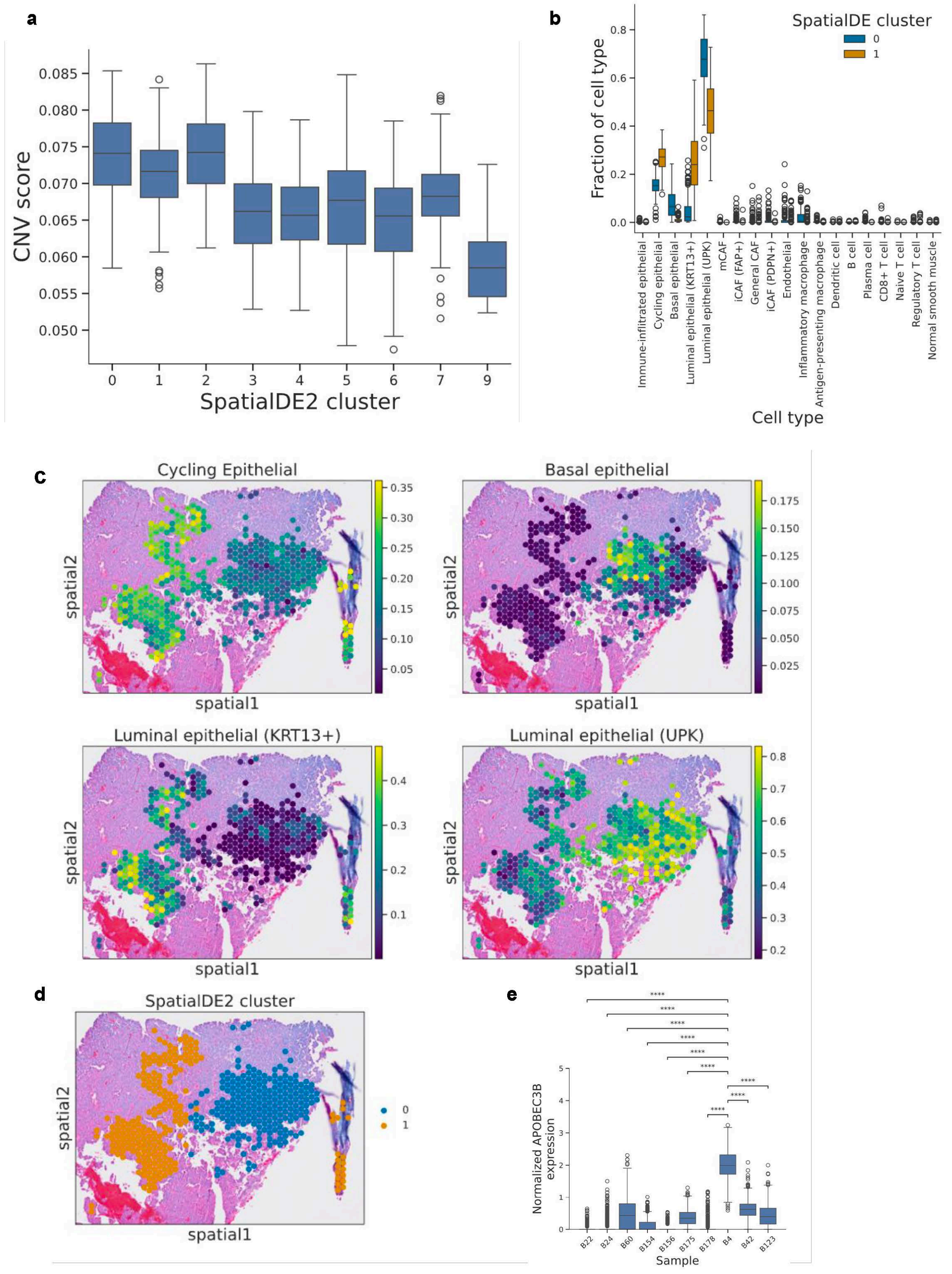


**Supplementary figure 12. ecDNA-enriched spatial clusters show different cell type composition in sample B4**. **a)** CNV score per spot for each SpatialDE2 cluster in sample B4. **b)** Comparison in the abundance of cell types between ecDNA-enriched clusters 0 and 1. **c)** Fraction of epithelial subtypes per spot for spatial clusters 0 and 1 in space. **d)** Spatial clusters 0 and 1 within the tissue architecture. **e)** Normalized expression of *APOBEC3B* per spot for all 10 Visium samples. A non-parametric Kruskal–Wallis test was used for statistical testing, followed by Dunn's test with Benjamini–Hochberg correction for multiple testing. Number of spots per sample: B22, n=504; B24, n=1171; B60, n=505;

B154, n=2525; B156, n=1826; B175, n=1102; B178, n=1256; B4, n=1284; B42, n=1037; B123, n=1507.

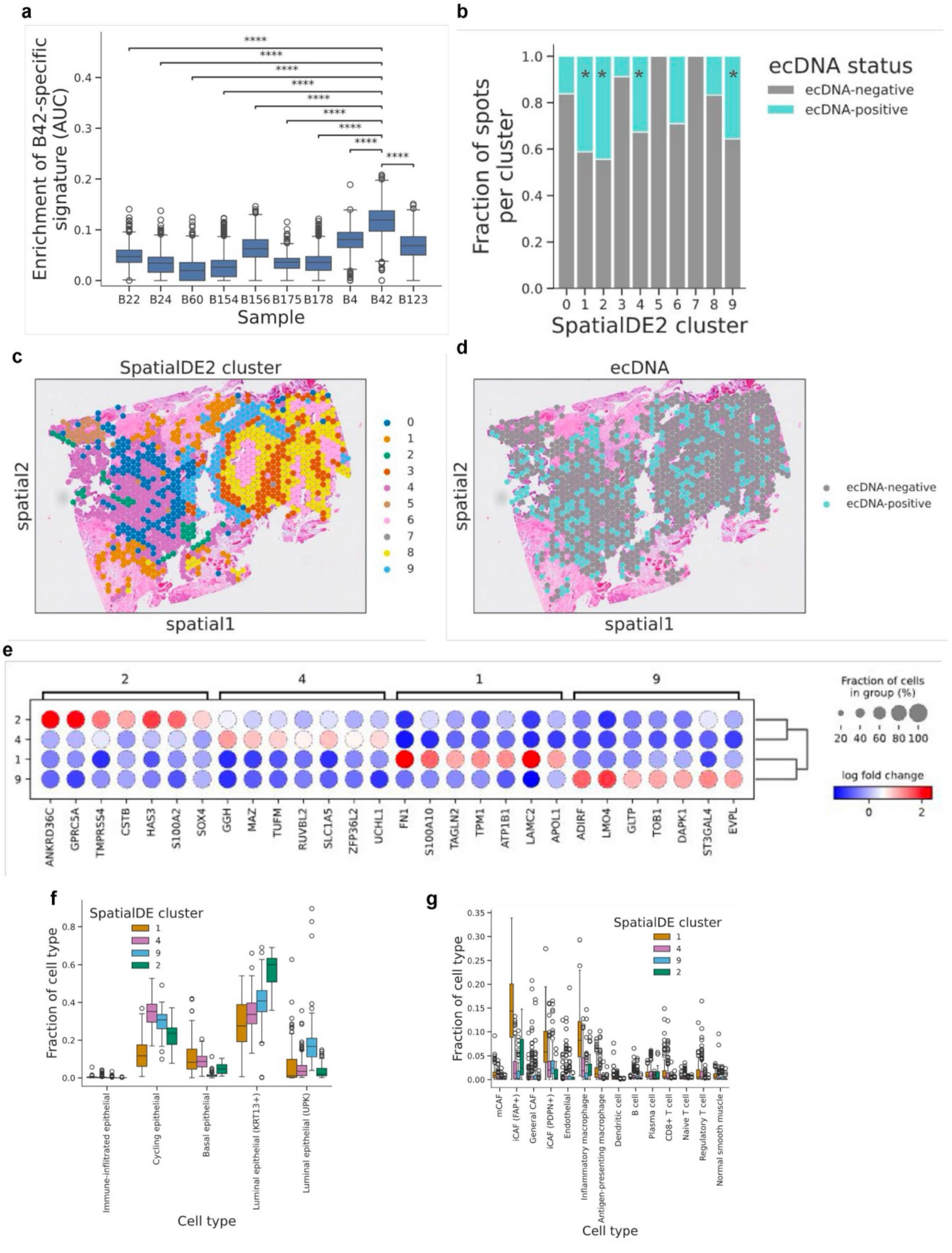


**Supplementary figure 13. Detection of ecDNA in Visium data for the sample B42. a)** AUC scores (enrichment) of the gene set consisting of genes on ecDNA in tumor B42, computed for each spot in all 10 Visium samples. A non-parametric Kruskal–Wallis test was used for statistical testing, followed by Dunn's test with Benjamini–Hochberg correction for multiple testing. Number of aneuploid spots per sample: B22, n=504; B24, n=1171; B60, n=505; B154, n=2525; B156, n=1826; B175, n=1102; B178, n=1256; B4, n=1284; B42, n=1037; B123, n=1507. **b)** Fraction of ecDNA-positive spots per

SpatialDE2 cluster from **c**. Stars annotate the clusters significantly enriched with ecDNA-positive spots (one-sided Fisher's exact test with Benjamini-Hochberg correction for multiple testing). **c)** Transcriptionally similar clusters in space computed by SpatialDE2. **d)** Predicted ecDNA status in space. **e)** Top 7 differentially expressed genes for all ecDNA-enriched clusters. **f)** Abundance of epithelial cell subtypes for ecDNA-enriched clusters 1, 2, 4 and 9. **g)** Abundance of non-epithelial cell types for ecDNA-enriched clusters 1, 2, 4 and 9.

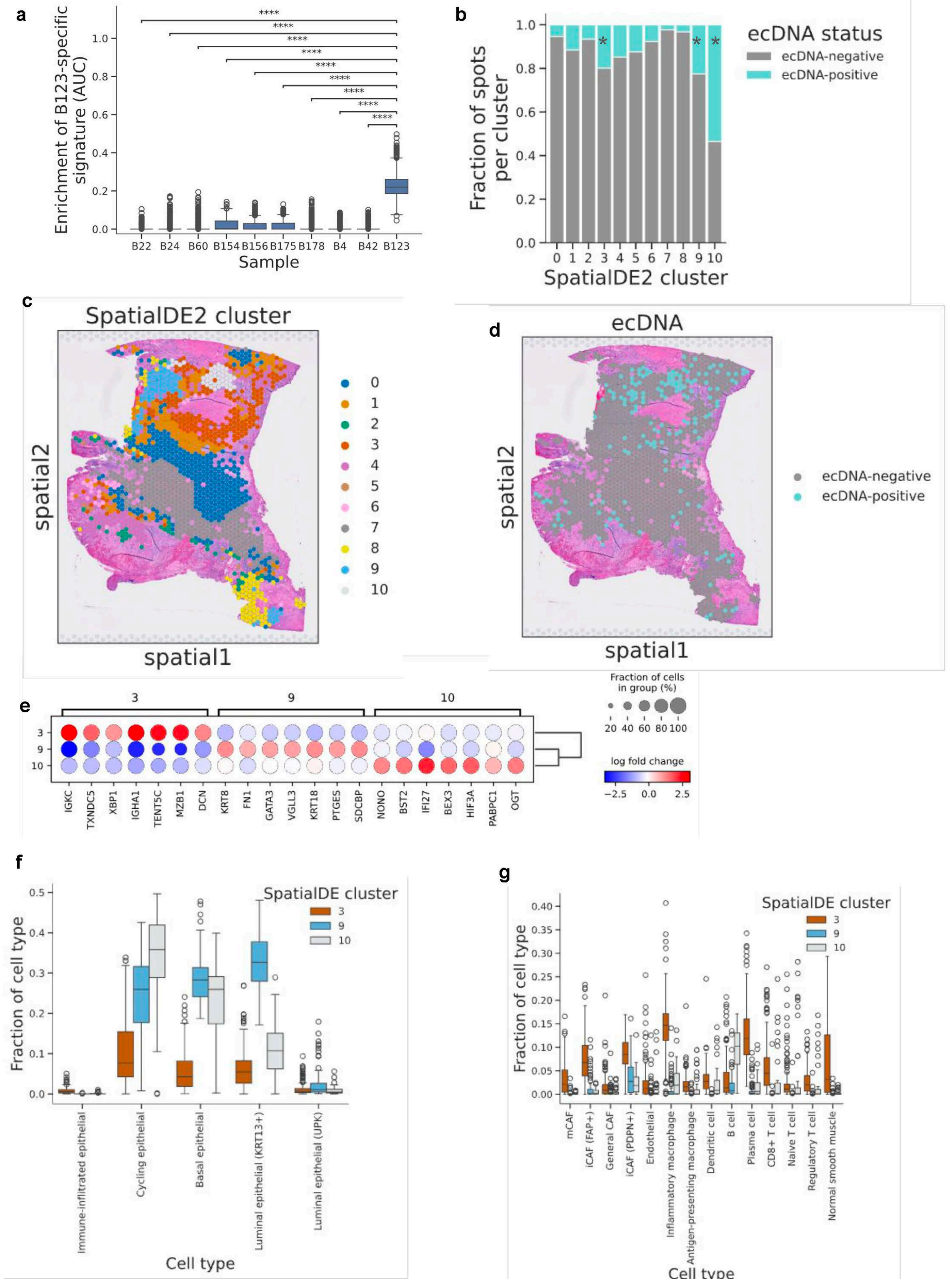
a
Enrichment of B123-specific signature (AUC)
Sample
B22 B24 B60 B154 B156 B175 B178 B4 B42 B123
****
b
Fraction of spots per cluster
SpatialDE2 cluster
ecDNA status
ecDNA-negative
ecDNA-positive
c
SpatialDE2 cluster
spatial1
spatial2
d
ecDNA
ecDNA-negative
ecDNA-positive
e
Fraction of cells in group (%)
log fold change
IGKC TXNDC5 XBP1 IGHA1 TENT5C MZB1 DCN KRT8 FN1 GATA3 VGLL3 KRT18 PTGES SDCBP NONO BST2 IFI27 BEX3 HIF3A PABPC1 OGT
f
Fraction of cell type
Cell type
SpatialDE cluster
Immune-infiltrated epithelial
Cycling epithelial
Basal epithelial
Luminal epithelial (KRT13+)
Luminal epithelial (UPK)
g
mCAF
iCAF (FAP+)
General CAF
iCAF (PDPN+)
Endothelial
Inflammatory macrophage
Antigen-presenting macrophage
Dendritic cell
B cell
Plasma cell
CD8+ T cell
Naive T cell
Regulatory T cell
Normal smooth muscle

**Supplementary figure 14. Detection of ecDNA in Visium data for the sample B123. a)** AUC scores (enrichment) of the gene set consisting of genes on ecDNA in tumor B123, computed for each spot in all 10 Visium samples. A non-parametric Kruskal–Wallis test was used for statistical testing, followed by Dunn's test with Benjamini–Hochberg correction for multiple testing. Number of aneuploid spots per sample: B22, n=504; B24, n=1171; B60, n=505; B154, n=2525; B156, n=1826; B175, n=1102; B178, n=1256; B4, n=1284; B42, n=1037; B123, n=1507. **b)** Fraction of ecDNA-positive spots per SpatialDE2 cluster from **c**. Stars annotate the clusters significantly enriched with ecDNA-positive spots (one-sided Fisher's exact test with Benjamini-Hochberg correction for multiple testing). **c)** Transcriptionally similar clusters in space computed by SpatialDE2. **d)** Predicted ecDNA status in space. **e)** Top 7 differentially expressed genes for all ecDNA-enriched clusters. **f)** Abundance of epithelial cell subtypes for ecDNA-enriched clusters 3, 9 and 10. **g)** Abundance of non-epithelial cell types for ecDNA-enriched clusters 3, 9 and 10.

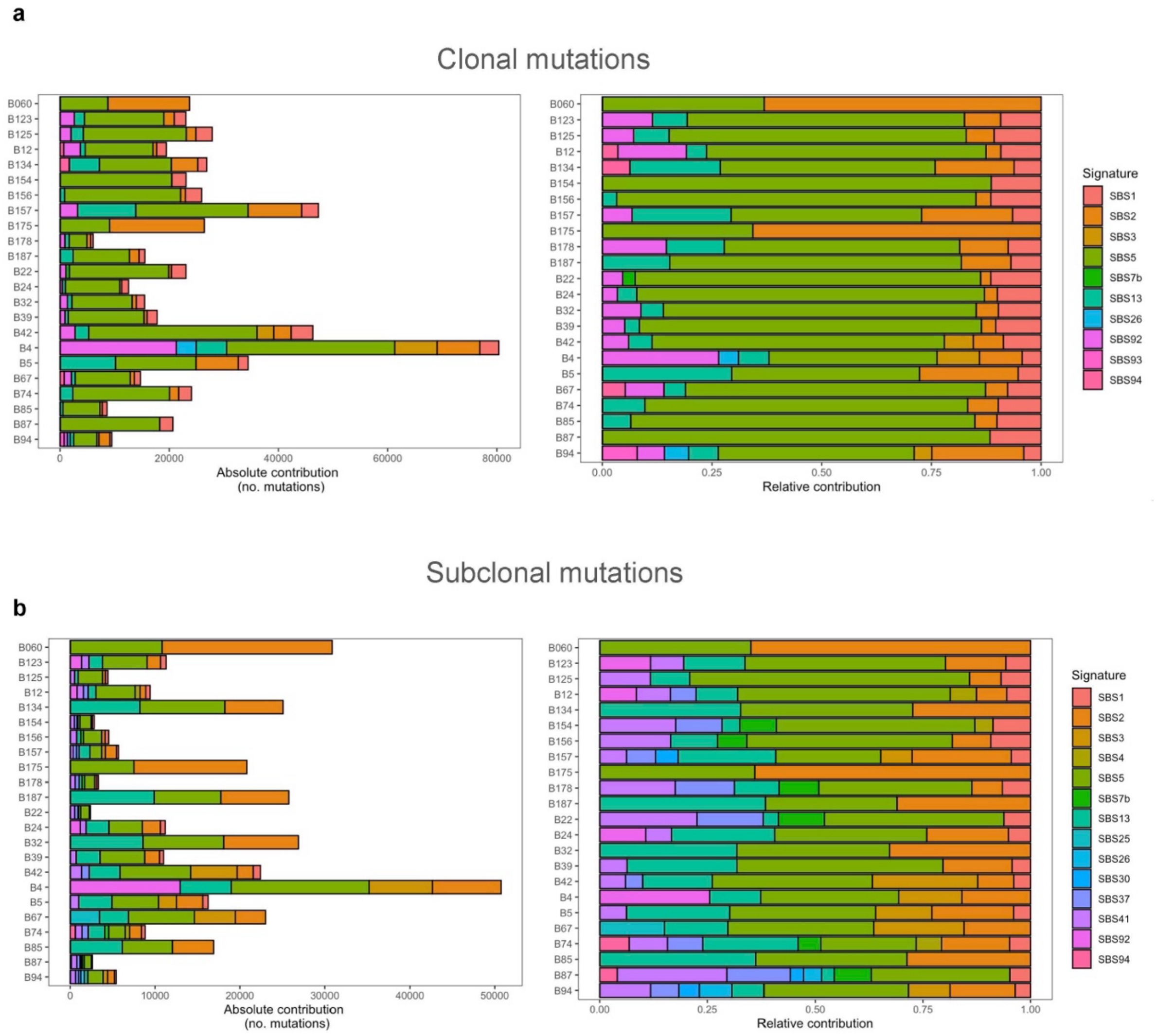


**Supplementary figure 15. a)** Relative and absolute contribution of each COSMIC signature in the patient cohort for clonal mutations. **b)** Relative and absolute contribution of each COSMIC signature in the patient cohort for subclonal mutations.

SUPPLEMENTARY TEXT

Germline variants in patient cohort

In the germline, we identified **12 high-impact pathogenic variants** in the following genes: *ARID4A*, *B3GLCT*, *DHTKD1*, *ENAM*, *FLG*, *GJB2*, *HMOX1*, *IQCB1*, *MSR1*, *NBN*, *RAB27A*, and *TSHB*.

Additionally, we detected **33 moderate-impact variants** across the following genes: *ABCA4*, *ABHD12*, *AGBL1*, *AGXT*, *BCO1*, *CD3G*, *CFHR5*, *CNGA3*, *CRYAA*, *MCM2*, *MLH3*, *MPEG1*, *MPL*, *MSH3*, *MYPN*, *NR2E3*, *NUBPL*, *OCA2*, *ORC4*, *PADI3*, *PCCB*, *PKHD1*, *PMP22*, *PRICKLE1*, *PRKDC*, *PRKN*, *PYGL*, *SERPINA6*, *SUCLG1*, *TYR*, *USH2A*, *WFS1*, and *XYLT1*.

Finally, we observed **two modifier variants** in *DTX4* and *TMEM107*.
